# Geographic divergence and the genomic basis of reproductive diapause in *Drosophila triauraria*

**DOI:** 10.1101/2025.09.19.677350

**Authors:** Takako Fujichika, Yuki Aoyama, Moe Onuma, Masafumi Nozawa, Aya Takahashi

## Abstract

Adjusting reproduction timing to environmental cues is essential for lifetime fitness. In many insects, reproductive diapause shows clinal variation along environmental gradients such as photoperiod and temperature. How such continuous trait variation may be encoded at the molecular level and maintained in the presence of gene flow remains largely elusive. The fruit fly *Drosophila triauraria* is distributed across a wide latitudinal range of the Japanese archipelago. Northern strains exhibit a strong photoperiodic reproductive diapause in females, whereas southern strains fail to arrest ovarian development even under short-day conditions at low temperatures. These distinct phenotypes and the presumable clinal variation in between, provide an ideal opportunity to examine the molecular basis of latitudinal divergence. We first investigated diapause induction in both females and males from previously reported and newly tested strains collected from the regions spanning ∼26–43°N along the Japanese archipelago. The assessment revealed continuous geographic variation in sensitivity to photoperiod and temperature. We then analyzed the whole-genome sequences of 21 strains, including 14 newly sequenced, to identify genomic regions underlying this divergence. In addition to the conventional F_ST_ analysis, we applied a “monophyletic window” approach suitable for limited sample sizes. The analysis identified a candidate region containing putative E-box and TER-box sequence motifs of the *timeless* (*tim*) gene, which has been previously implicated in diapause regulation in multiple insect species. The quantitative PCR analysis further supported a partial association between the *tim* expression and the incidence of female diapause. These findings reinforce the growing evidence for a role of circadian clock genes in the adaptive regulation of reproductive diapause and demonstrate the utility of tree-based approaches for detecting genomic regions of geographic divergence.

## INTRODUCTION

Adjusting internal physiological states to environmental fluctuations is essential for survival and reproductive success. Traits related to the sensitivity to environmental cues or the magnitude of physiological responses often maintain polymorphisms due to fluctuating or unpredictable fitness outcomes (e.g., Johnson et al., 2023). Such traits may also exhibit intraspecific clinal variation when key environmental factors follow a continuous geographical gradient (reviewed in Hut et al., 2013; Mayekar et al., 2022). Sensitivity to photoperiod and temperature cues that induce diapause in insects is a well-documented example (reviewed in Bradshaw & Holzapfel, 2007; Denlinger, 2023). Despite decades of studies using various insect species, the molecular mechanisms that enable continuous variation in diapause induction along clinal environmental gradients, and the processes that maintain such variation in the presence of gene flow, remain elusive.

Many studies have attempted to identify loci involved in the photoperiodic regulation of diapause across clinal populations to tackle this question. In *Drosophila melanogaster*, specific genes with large effects have been identified, such as *ls*-*tim* in European populations (Tauber et al., 2007) and *couch potato* in North America (Schmitt et al., 2008). However, these associations appear inconsistent across different geographic regions (Lee et al., 2011; Zonato et al., 2016; 2018), suggesting a polygenic nature of the phenotype (Bradshaw et al., 2012; Laruson et al., 2020; Barghi et al., 2020). Recent genomic resequencing studies in Lepidoptera, comparing population differentiation or conducting large-scale phenotype-genotype association analyses, have identified additional candidate genes, many of which are related to circadian regulations (Kozak et al., 2019; Pruisscher et al., 2021; Lindestad et al., 2022; Yu et al., 2023; Zheng et al., 2025). Although single genetic variants rarely explain the full extent of phenotypic variation in many cases (e.g., Pruisscher et al., 2018; Lindestad et al., 2024), the association between the circadian clock and photoperiodic regulation is widely recognized Lepidoptera as well as in other insect species (Ikeno et al. 2010; Chang & Meuti, 2020; Hasebe & Shiga, 2025). Whether this association represents a consistent pattern across diverse insect species requires further investigation, as it has not been detected in some studies (e.g., Emerson et al., 2009; Lirakis et al., 2022). At the same time, despite advances in genomic technologies, collecting sufficiently large sample sizes across broad geographic ranges, alongside phenotypic assays and genome sequencing, continues to be a significant challenge.

*Drosophila* species, many of which likely to have originated in tropical regions, have expanded to occupy habitats spanning a wide latitudinal range (Throckmorton, 1975; Lemeunier et al., 1986; Markow & O’Grady, 2006). Their migratory ability has also facilitated the expansion of habitats within species (e.g., Sprengelmeyer et al., 2020), providing opportunities to investigate both shared and convergent genetic bases of clinal variation. Because of the direct fitness consequences, clinal variation in reproductive diapause, in which reproduction is arrested under unfavorable environmental conditions, is documented in multiple *Drosophila* species inhabiting temperate to subarctic regions. The diapause incidence increases with latitude in many of these species, reflecting local adaptation despite ongoing gene flow. For example, diapause incidence varies with latitude between ∼35–70°N in *D. montana* (Tyukmaeva et al., 2020; Lankinen et al., 2023) and *D. littoralis* (Lankinen, 1986), and between ∼34–43°N in *D. auraria* (Minami & Kimura, 1980; Kimura, 1984; Pittendrigh & Takamura, 1987) and *D. lacertosa* (Ichijo, 1986). Similarly, clinal variation of diapause incidence has been intensively studied in *D. melanogaster* along the eastern coast of North America (∼25–45°N) and Europe (e.g., Schmidt et al., 2007; Tauber et al., 2007).

In this study, we focus on *D. triauraria*, a member of the auraria complex, which inhabits wide latitudinal ranges (∼26–43°N) across the Japanese archipelago (Kimura, 1987). In this species, a strong photoperiodic reproductive diapause has been reported in northern strains, whereas, notably, southern strains fail to arrest ovarian development even under short-day conditions at 15°C (Kimura, 1983; 1988; Yamada & Yamamoto, 2011). These distinct phenotypes, together with the presumable clinal variation in between, provide an ideal opportunity to examine the molecular basis of latitudinal divergence.

Here, we investigate the sensitivity to temperature and photoperiod in reproductive diapause induction in both females and males of *D. triauraria* from mid-latitude regions alongside previously reported and newly tested strains from high- and low-latitude populations. We then examined the degree of divergence from the distribution of a previously described polymorphism in genital morphology (Onuma et al., 2022), and also by a phylogenetic analysis using the whole-genome sequences of the 21 *D. triauraria* strains. Next, to finding signatures of regional divergence, we applied a “monophyletic window” approach, in addition to the conventional F_ST_ analysis. A candidate region contained the *timeless* (*tim*) gene; thus, we conducted quantitative PCR analysis to analyze the association between the *tim* expression and the female reproductive diapause incidence. We discuss the involvement of a circadian clock gene in the adaptive regulation of reproductive diapause in *D. triauraria* and demonstrate the utility of the monophyletic window approach for identifying genomic regions showing geographic divergence.

## MATERIALS AND METHODS

### Fly strains

The *Drosophila* strains used in this study are described in Table 1. The sampled locations on the map are shown in Figure 1. All strains were maintained at 20°C under the 15 h light: 9 h dark cycle. All flies were fed with standard corn meal food mixed with yeast, glucose, and agar.

**FIGURE 1.**
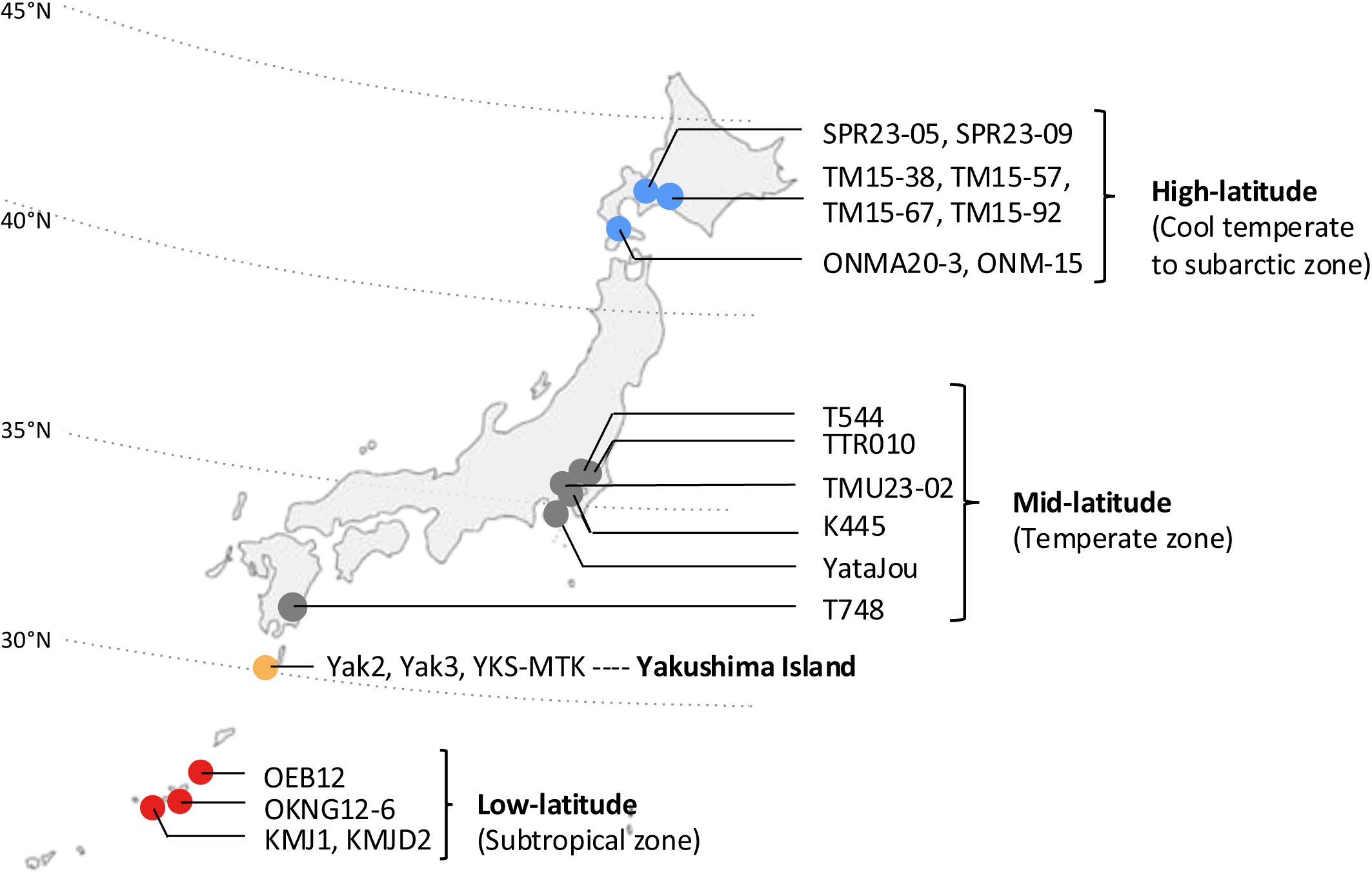
Collection sites of *D. triauraria* strains from Japanese archipelago used in this study. A total of 21 isofemale lines of *D. triauraria* that were established from natural populations were used. Circles indicate collection sites: blue, gray, and red denote high-, mid-, and low-latitude regions, respectively, while orange represents Yakushima Island, which has a unique mixed climate distinct from the other regions.

**Table 1.**
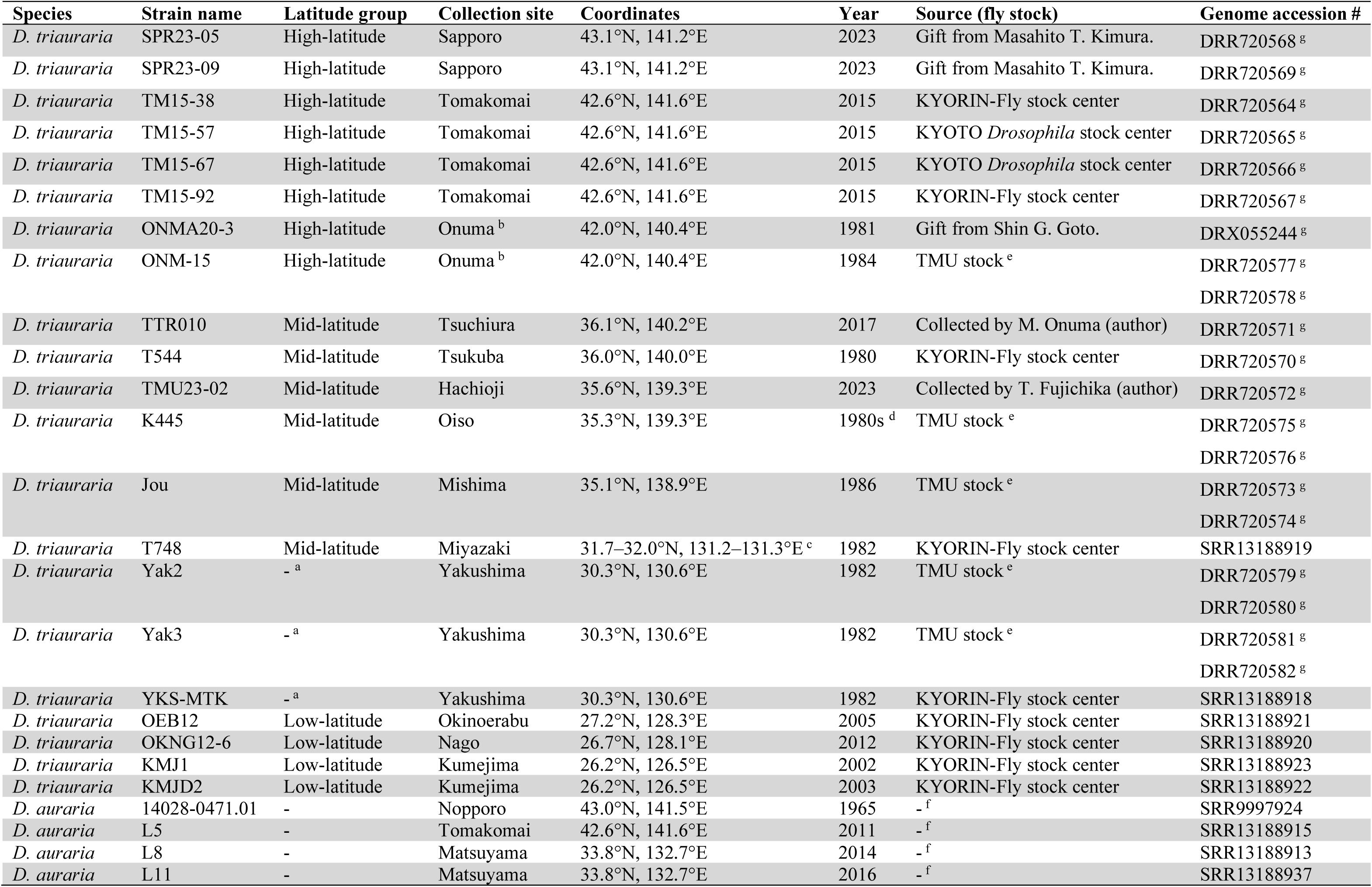
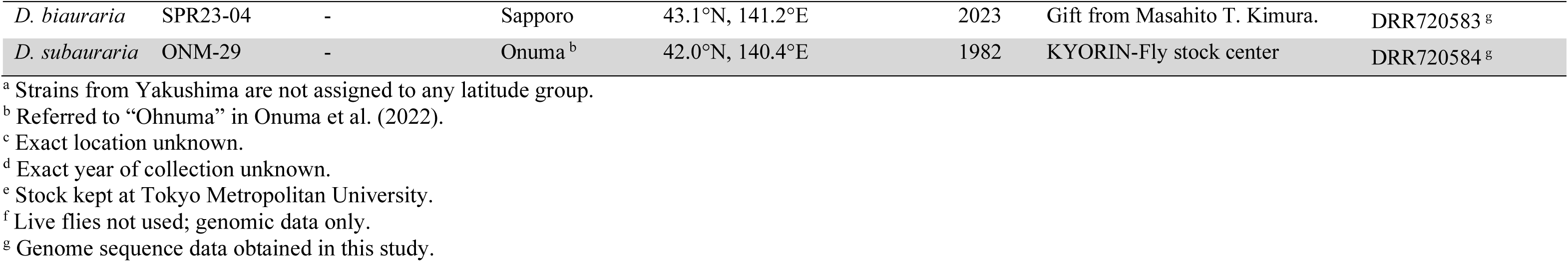
Strains of *D. triauraria*, *D. auraria*, *D. biauraria*, and *D. subauraria* used in this study.

### Female diapause assay

The flies were not exposed to the test conditions prior to eclosion, as exposure to low temperatures during embryonic and larval development results in high mortality. Ten virgin females collected within 6 h of eclosion were placed into the food vials and reared under an 8 h light: 16 h dark (8L:16D) or a 16 h light: 8 h dark (16L:8D) cycle, at 12°C, 15°C, and 18°C. The 15°C and 18°C conditions were adopted from the previous studies of this species (Kimura, 1983; Yamada & Yamamoto, 2011), and the 12°C condition was introduced in addition to those temperatures. The 16L:8D and 8L:16D testing light cycles were set to be slightly longer and shorter, respectively, than the natural day lengths (hours between sunrise and sunset) at the summer peak (∼15.5 h) and winter peak (∼9 h) at the collection sites of the high-latitude strains. Flies were dissected after 21 days at 12°C, 16 days at 15°C, and 14 days at 18°C, according to the timing of sexual maturation inferred from the previous studies (Kimura, 1983; Oguma et al., 1996). The females were dissected in phosphate-buffered saline (PBS), and the ovary images were taken by a CCD camera (DP73, Evident) attached to a microscope (SZX16, Evident). The developmental stages of eggs inside the ovarioles were determined according to King (1970). Diapaused ovaries were defined as those with egg stages 0 to 7 (Figure 2).

**FIGURE 2.**
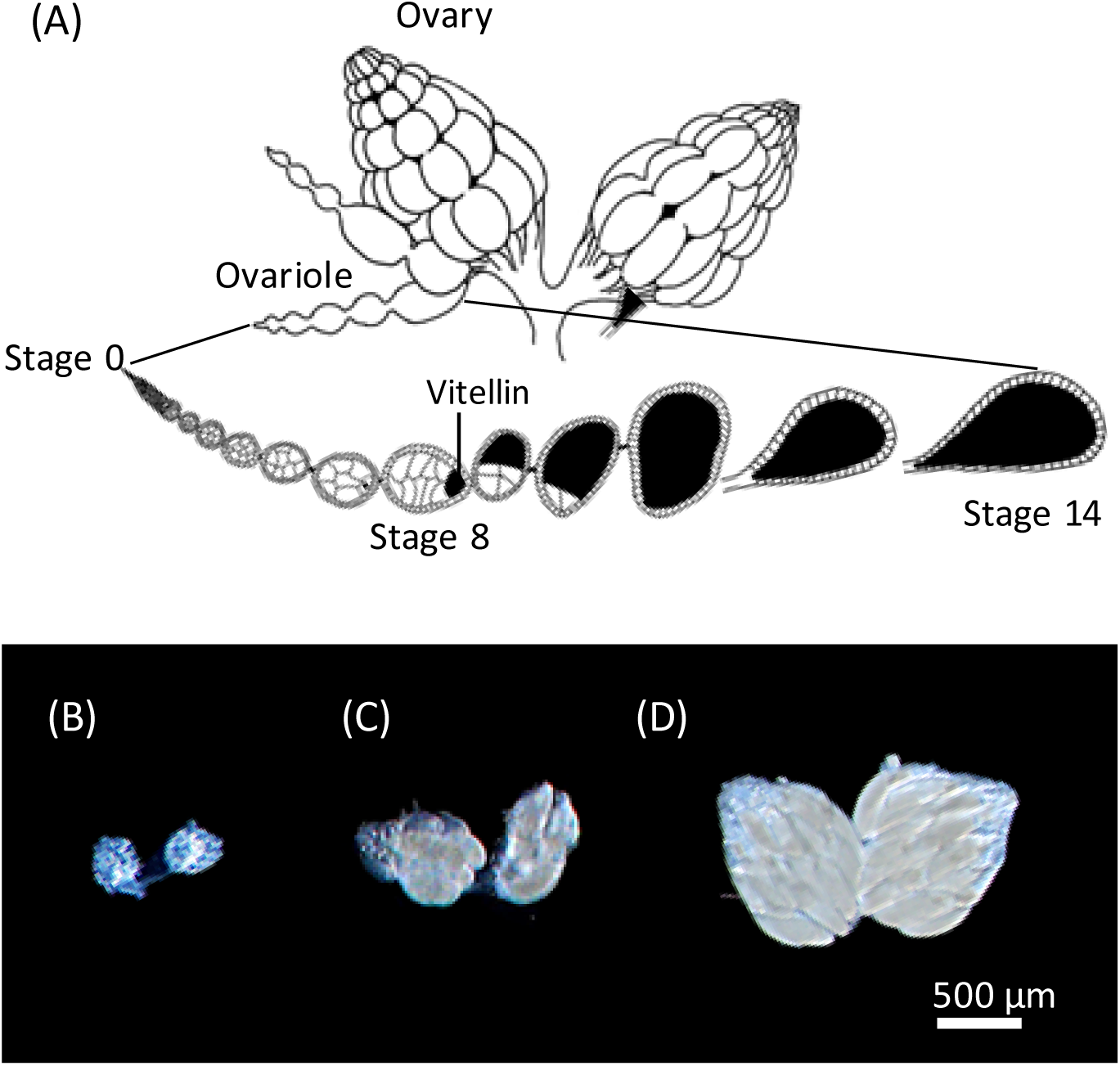
Reproductively diapaused and non-diapaused ovaries of *D. triauraria*. (A) Schematic illustration of a pair of *Drosophila* ovaries, modified from King (1970). Each ovary consists of multiple ovarioles, where oogenesis progresses from stage 0 to 14 (mature egg). (B) Diapaused ovaries, lacking eggs at ≥ stage 8 (vitellogenic stages). (C) Non-diapaused ovary with nine eggs at ≥ stage 8. (D) Fully developed ovary.

### Measurements of Accessory gland (AG)

Ten non-mated males collected within 6 h of eclosion were dissected immediately or placed into the food vials and reared under an 8L:16D or 16L:8D light/dark cycle at 12°C for 21 days. The males were dissected in PBS and their accessory glands were fixed in 4% paraformaldehyde (PFA) in PBS for 30 min, followed by three washes in PBST (0.05% Tween 20). The samples were then transferred into PBS. For imaging, 7 μL VectorShield with DAPI (Vector Laboratories) was used for mounting on the slides. Images were taken by a CCD camera (DP73, Evident) attached to an inverted microscope (IX73, Evident). The obtained images were analyzed for area measurement using ImageJ (Schneider et al. 2012).

### Male genitalia dissection

Genitalia from 10 males aged 7–10 days were dissected in glycerol after boiling whole bodies in 10% potassium hydroxide at 95°C for more than 10 min to digest soft internal tissues and facilitate clearing. Dissections were performed under a stereomicroscope (SZX7, Evident). Images were acquired using an inverted microscope (IX73, Evident) equipped with a CCD camera (DP73, Evident). Image stacks were processed with CellSens software (Evident) to generate depth-composited TIFF files. Terminology and identification of phallic structures followed Rice et al. (2019) and Onuma et al. (2022) for *D. triauraria*.

### Genome sequencing

Genomic DNA was extracted from 40 individuals (20 males and 20 females) per strain for the nine *Drosophila triauraria* strains (SPR23-05, SPR23-09, TM15-38, TM15-57, TM15-67, TM15-92, T544, TTR010, and TMU23-02), along with one *D. biauraria* strain (SPR23-04) and one *D. subauraria* strain (ONM29). DNA extraction was performed using the QIAGEN Genomic-tip 20/G kit (QIAGEN), following the protocol described by Solares et al. (2018). DNA concentrations were quantified using a Qubit dsDNA Quantification Assay Kit (Invitrogen). For each sample, 1 µg of DNA was sent to Filgen, Inc. for library construction and sequencing using a NovaSeq6000 platform with 150 bp paired-end reads.

Additionally, genomic DNA was extracted separately from 10 females and 10 males from each of the five *D. triauraria* strains (ONM-15, K445, Jou, Yak2, and Yak3) using the Geno Plus Genomic DNA Extraction Miniprep System (Viogen). DNA samples (50–250 ng per sample) were used for library preparation with the Collibri PCR-free ES DNA Library Prep Kit for Illumina (Invitrogen). Quality assessment of the libraries was performed using an Agilent 2100 Bioanalyzer (Agilent Technologies). The libraries were then sent to Macrogen Inc., Korea, for sequencing by using a HiseqX with 150 bp paired-end reads. Read depth and quality information for each strain are listed in Table S1.

### Quality control and variant calling of the genomic sequence data

Quality control of the sequencing reads was performed using FastQC (Andrews, 2010). Adapter trimming and quality filtering (QC > 30) were conducted using Trim Galore! (Krueger, 2021). Reads were then mapped to the *D. triauraria* reference genome (GCA_014170255.2, RU_Dtri_1.1, NCBI) using Bowtie2 (Langmead & Salzberg, 2012). The mapped reads were sorted using SAMtools (Li et al., 2009), and variant calling was performed using BCFtools v1.18 (Danecek et al., 2021). The quality of the variant calling process was assessed using MultiQC (Ewels et al., 2016) based on the BCFtools v1.18 output. Post-calling variant filtration was performed using BCFtools v1.18. Variants with QUAL ≤ 10, DP > 120, or MQBZ < −3 were filtered out, and the remaining variants were output in variant call format (VCF).

### Maximum likelihood tree construction

A FASTA file for each strain was constructed by replacing the reference sequence with the single-nucleotide polymorphisms and deletions (replaced by "-") in the variant call format (VCF) file. The FASTA files from all strains were merged (alignment kept from the original VCF file) and converted to PHYLIP format. Maximum likelihood phylogenetic trees were constructed using IQ-TREE (Minh et al., 2013; 2020; Nguyen et al., 2015) with ultrafast bootstrap (UFBoot) analysis (1,000 replicates) and the GTR+G+I substitution model. Gaps were removed prior to running IQ-TREE. To construct trees from specific 5 kb windows, the focal regions were extracted from the merged FASTA file using SeqKit2 (Shen et al., 2024), in prior to PHYLIP conversion.

### Monophyletic window analysis by maximum parsimony tree construction

To conduct a genome-wide sliding window analysis, VCF files from all strains were directly merged (without first constructing FASTA files). The merged VCF file was then divided into 5 kb windows with a 1 kb step size using BCFtools v1.18, and converted to PHYLIP format using vcf2phylip.py (Ortiz, 2019). MPBoot v1.1 (Hoang et al., 2018) was used to generate maximum parsimony phylogenetic trees from each window with 1,000 UFBoot replicates. Monophyly of a particular latitudinal group of strains was analyzed using ape 5.0 (Paradis & Schliep, 2019) and phangorn (Schliep, 2011) packages in R (R core team, 2025). Windows with trees containing monophyletic inner branches with UFBoot values ≥ 90 were considered as candidate genomic regions. The phylogenetic trees were visualized and examined using FigTree v1.4.4 (Rambaut, 2018). Other window sizes (10, 20, and 100 kb) were also analyzed for comparisons. Smaller window sizes provide higher resolution but reduced statistical power due to the limited number of variants. The window size of 5 kb was selected as the smallest size for which more than 90% of windows contained at least 100 variants. To identify genes within the candidate regions, a local BLASTN search (Altschul et al., 1990) using BLAST v2.15.0 (Camacho et al., 2009) was conducted against the *D. melanogaster* transcripts (GCF_000001215.4_Release_6_plus_ISO1_MT_rna.fna.gz), retrieving the hits with E-values < 0.01.

### Genetic differentiation analysis

To assess genetic differentiation between groups, Weir and Cockerham’s F_ST_ value (Weir and Cockerham, 1984) was calculated using VCFtools (Danecek et al., 2011) for each 5 kb window. The weighted F_ST_ statistic was used to account for varying sample sizes. The results were visualized in R using the ggplot2 package (Wickham, 2016).

### Sanger sequencing of the two adjacent monophyletic windows (6 kb) including *tim*

Genomic DNA was extracted from a single female using the NucleoSpin DNA Extraction Kit (Macherey-Nagel). PCR amplification of the 6 kb region was performed using KAPA HiFi HotStart Ready Mix (KAPA Biosystems) or Takara Ex Premier DNA Polymerase (Takara Bio). Two sets of primer pairs were used: 1) 5’-CAGTGCTGATAAACCCCCTTACCTA- 3’ or 5’-CACTTGGTTATGAAAGCCCTAACCT- 3’ and 5’-GCCAGTTGTAGGTACTTCTCCTCCT- 3’; 2) 5’-AAATACCGGTTATGGACTGGTTACTAGC- 3’ and 5’- AAGTCCTCCAAGAGGATGCCATAATAGT- 3’. PCR products were purified using AMPure XP beads (Beckman Coulter) or QIAquick Gel Extraction Kit (QIAGEN). Sanger sequencing was performed using BigDye Terminator v3.1 Cycle Sequencing Kit (Applied Biosystems, Thermo Fisher Scientific) and ABI 3130xl Genetic Analyzer (Applied Biosystems, Thermo Fisher Scientific). Sequence assembly was conducted using BioEdit (Hall, 1999), and the resulting FASTA files were compiled and aligned using MEGA7 (Kumar et al., 2016). The heterozygotes were replaced by the reference sequence manually or using Mitsucal (Suzuki et al., 2018).

Transcript models of *tim* were built by aligning a publicly available ovary transcripts (SRR11780982) to the reference genome (GCA_014170255.2, RU_Dtri_1.1, NCBI) using HISAT2 (Kim et al., 2019). Adapter trimming and quality filtering (QC > 30) were conducted using Trim Galore! in prior to alignment. The aligned reads were sorted into BAM files with SAMtools and assembled into transcripts using StringTie (Pertea et al., 2015). Open reading frames (ORFs) were predicted with TransDecoder (Haas et al. 2013) with a minimum length of 50 amino acids and subsequently filtered by BLASTP against *D. melanogaster* protein sequences (FlyBase dmel-all-translation-r6.36.fasta). BLASTP hits were used to retain only ORFs with significant matches (E-value < 1e-10).

### 3’ Rapid amplification of cDNA ends (3’ RACE) of *tim* isoforms

One hundred heads from *D. triauraria* females reared at 20°C under a 15L:9D light/dark cycle were flash-frozen in liquid nitrogen at Zeitgeber-time (ZT)4, corresponding to 4 h after lights on. Total RNA was extracted using the PureLink RNA Mini Kit with TRIzol Reagent (Thermo Fisher Scientific). RNA concentration was measured using a NanoDrop 2000 spectrophotometer (Thermo Fisher Scientific). cDNA synthesis was performed using PrimeScript III Reverse Transcriptase (Takara Bio) with 1 μg of total RNA as a template. The oligo (dT) primer was used at a final concentration of 100 pmol/μL. PCR amplification was carried out using DNA Ex Premier Polymerase (Takara Bio) with 5’-GAGCTCAAGCTTAATCTGTTAAAAGAGTTTACAGTGG-3’ and 5’-CCAGTGAGCAGAGTGACGAGGACTCGAGCTCAAGCTTTTTTTTTTTTTTTTT-3’. PCR products were analyzed by agarose gel electrophoresis, and target bands (800–1,000 bp and 3,000–3500 bp) were excised and purified using NucleoSpin Gel and PCR Clean-up Kit (Takara Bio). The purified DNA fragments were cloned into the pBluescript II SK(+) vector (Agilent Technologies). The vector and PCR products were digested by NotI and treated with shrimp alkaline phosphatase (SAP) for dephosphorylation, followed by ligation using T4 DNA Ligase (Takara Bio). The ligated plasmids were transformed into competent *E. coli* cells. Plasmid DNA was extracted from colonies containing plasmids with a PCR product insertion using Cica Geneus Plasmid Kit (Kanto Chemical). The inserted PCR products were then subjected to Sanger sequencing as described above.

### Quantification of clock genes or *tim* isoforms transcription level

Ten virgin females of ONMA20-3 were collected within 6 h of eclosion and reared in food vials (10 individuals per vial) under an 8L:16D or 16L:8D light/dark cycle at 12°C. After 7 days, the females were immediately frozen in liquid nitrogen every 4 h and stored at −80°C. The same protocol was applied to 15 additional strains, except that the samples were collected at 8 h and 16 h after lights on under the respective 8L:16D or 16L:8D light/dark cycle.

To extract RNA, 10 frozen heads per sample were detached by vortexing for 5 seconds and placed into 2 mL microtubes containing 400 µL TRIzol Reagent and 1.2 mm (diameter) zirconia-silica beads. Tubes were shaken at 3,200 rpm for 2 min using a Beads Crusher µT-12 (Taitec). Chloroform (160 µL) was added, and total RNA was purified using the silica-gel-based method of Boom et al. (1990). First-strand cDNA was synthesized from 70 ng total RNA using the PrimeScript RT Reagent Kit with gDNA Eraser (Takara Bio).

Realtime reverse transcription quantitative PCR (RT-qPCR) was conducted using TBGreen Premix Ex Taq II (Tli RNaseH Plus, Takara Bio) in a total reaction volume of 12.5 µL. Each reaction contained 6.25 µL of TBGreen Premix, 2 µL of the 3-fold diluted cDNA sample, and 10 pmol each of forward and reverse primers (*tim*-*L*: 5’-CCCGCATGTATGTAAGCGATG- 3’ and 5’-AACTCGGATCTGGGATCTGCT- 3’, *tim*-*sc*: 5’- TGAACACTCCTCCAAAGTCACC- 3’ and 5’-GCATCTTAATCCTTACCAAATGG- 3’). Reactions were performed using the Dice Real Time System II thermal cycler (Takara Bio). The standard curves generated from 5-fold serial dilutions of an arbitrarily selected sample (the third replicate of the OEB12 samples collected at ZT16 under 16L:8D light/dark cycle) were used. Each sample was analyzed in technical duplicate, with three biological replicates for each condition.

## RESULTS

### Geographic variation in female diapause induction

Previous studies have reported a strong photoperiodic induction of reproductive diapause in females of *D. triauraria* from subarctic regions (42–44°N) and the contrasting non-diapausing phenotype of those from subtropical regions (26–28°N) of the Japanese archipelago (Kimura, 1983; 1988; Kimura & Yoshida, 1995; Yamada & Yamamoto, 2011). Strains from the template regions in between exhibited various levels of photoperiodic induction of reproductive diapause at 15°C (Kimura, 1983; 1988). To investigate the geographic variation in photoperiodic responses across temperatures, we measured the proportion of ovaries in reproductive diapause at 12°C, 15°C, and 18°C under either an 8L:16D or 16L:8D light/dark cycle using 21 strains from high- to low-latitude localities (Table 1, Figure 3).

**FIGURE 3.**
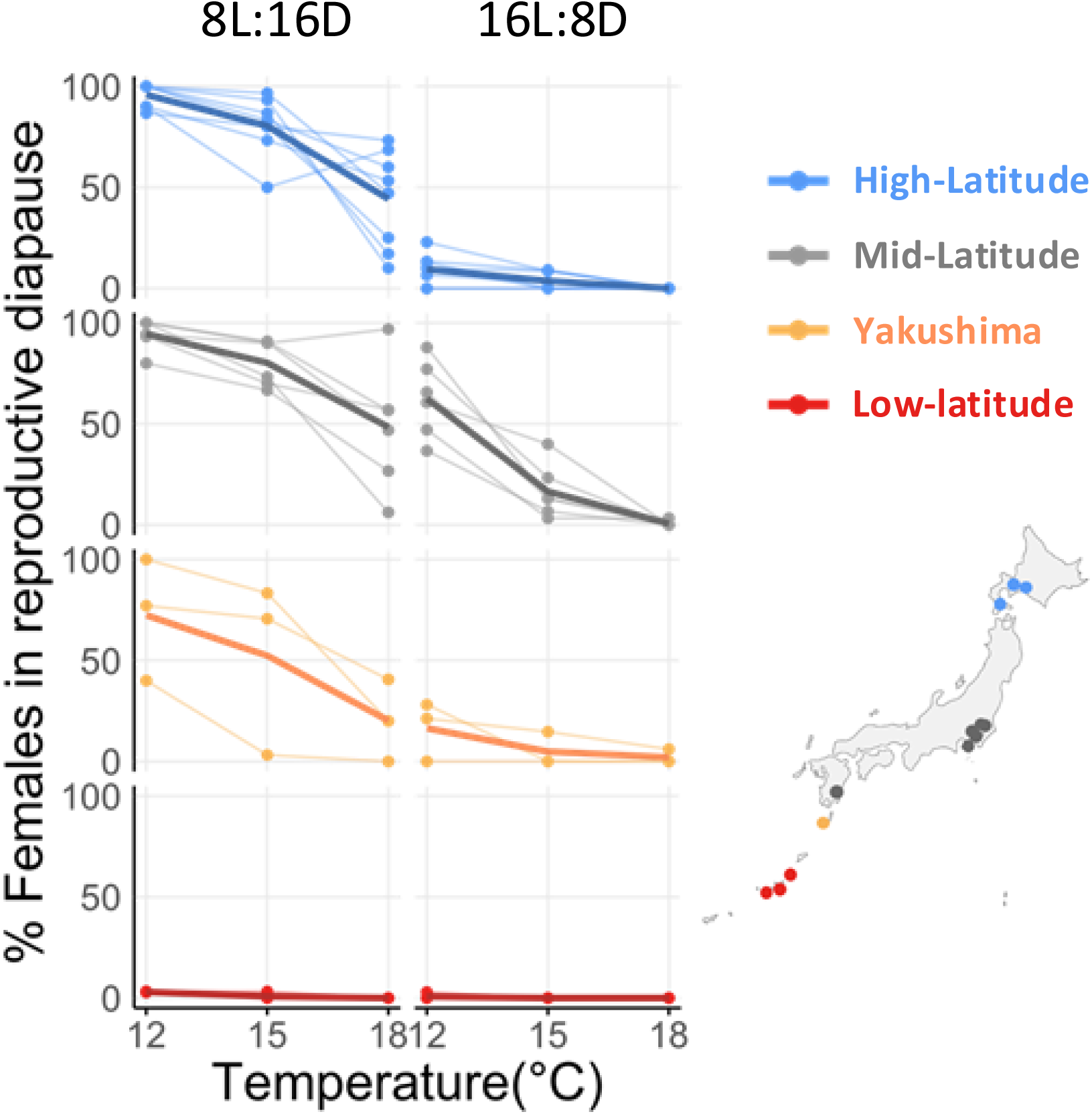
Effects of temperature and photoperiod on reproductive diapause incidence across regional populations of *D. triauraria*. The proportion (%) of female individuals in reproductive diapause under different temperatures (12°C, 15°C, and 18°C) and photoperiods (8L:16D or 16L:8D light/dark cycle). The colors indicate different latitudinal regions: blue, grey, orange, and red indicate high-latitude, mid-latitude, Yakushima, and low-latitude regions, respectively. Thick solid line connects the mean at each temperature.

Consistent with the previous studies, the strains from low-latitude regions showed almost no diapause incidence under any condition, showing continuous ovarian development. In contrast, strains from high-latitude regions induced diapause under short-day conditions in a temperature-dependent manner but went into reproductive mode under the long-day photoperiod regardless of temperature. Reproducing under long-day conditions at low temperature may be an adaptive strategy to cope with short summers in high-latitude localities.

Mid-latitude strains showed photoperiod-dependent diapause induction at 15°C and 18°C, like high-latitude strains. However, at 12°C, approximately half of the females induced reproductive diapause under the long-day condition. These results suggest that although mid-latitude strains respond to photoperiods like high-latitude strains, their photoperiodic response at low temperature is reduced, indicating a higher sensitivity to temperature. Three strains from Yakushima Island, which is known for its unique climate including subtropical in the coastal area to near subarctic in the mountainous ranges > 1,800 m, exhibited variable extent of diapause induction under all temperature conditions.

### Diapause induction in males

While reproductive diapause is more pronounced in females compared to males in *Drosophila* species, males also exhibit reduced reproductive activity under unfavorable conditions (Kimura, 1988; Kubrak et al., 2016; Ala-Honkola et al., 2020). The cost of reproduction typically differs between males and females (Bateman, 1948; Trivers, 1972); therefore, they may apply different criteria when allocating energy to reproduction. We investigated this possibility by examining male reproductive status under the 8L:16D or 16L:8D light/dark cycle at 12°C using 12 strains (four each from high-, mid-, and low-latitude populations). Yakushima strains were not included in the male accessory gland analysis because the diapause phenotypes of the females were variable and the variation was not associated with latitude, which is the primary focus of this study.

Reduction of testis and accessory gland size at low temperature and short photoperiod has been reported in *D. melanogaster* (Kubrak et al., 2016). However, unlike egg development in ovarioles, which has a checkpoint prior to the initiation of vitellogenesis, when environmental condition is assessed before investigating energy into egg production (Giorgi & Deri, 1976; Drummond-Barbosa & Spradling, 2001), there is no explicit point of arrest in male reproductive organs. Thus, we measured the size of accessory glands, where costly proteins are produced to be injected into the female reproductive tract during copulation.

The accessory gland sizes were measured from the males of all 12 strains kept 21 days after eclosion under the 8L:16D or 16L:8D light/dark cycle at 12°C (Figure 4A). For comparison, those of newly eclosed males within 6 h of eclosion were also measured and the 95% confidence interval (CI) was calculated from the measurement of all strains except T544 (Figure S1). Because the accessory glands of T544 males were significantly larger than those of Jou, OEB12, and OKNG12-6, the 95% CI was calculated separately for this strain. In this study, the males were conservatively classified as reproductively arrested if their accessory gland size was below the upper 95% CI limit of the newly eclosed males. Individuals with accessory gland sizes exceeding this threshold were classified as developing.

**FIGURE. 4.**
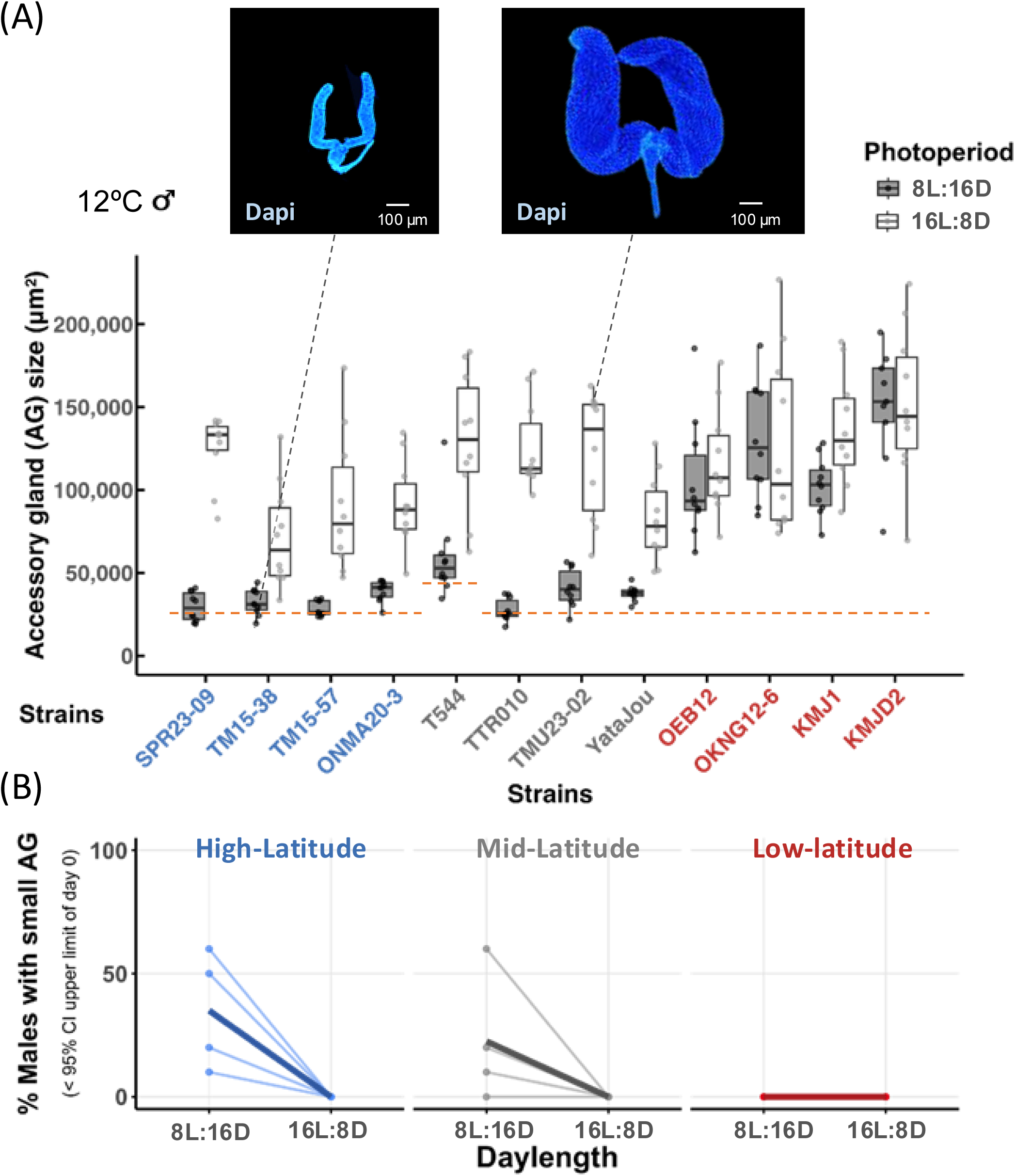
Effects of temperature and photoperiod on accessory gland (AG) growth across regional populations of *D. triauraria*. (A) Sizes of AGs after 21 days under different photoperiods (8L:16D or 16L:8D) at 12°C. Two representative images of AGs are shown above the graph. Orange dashed lines indicate the upper limit of the 95% CI calculated within strain for T544, and for the remaining strains, from the pooled data excluding T544. The separate calculation for T544 was performed due to its significant difference from other strains (see Figure S1). (B) Proportion (%) of males with AG sizes below the 95% CI upper limit of day 0 after 21 days under 8L:16D or 16L:8D at 12°C. Colors indicate latitudinal regions: blue, grey, and red indicate high-latitude, mid-latitude, and low-latitude regions, respectively. Thick solid line connects the means at each temperature.

The proportion of males under reproductive arrest are shown in Figure 4B. Unlike females, no male in any of the 12 strains met the criteria of reproductive arrest under the 16L:8D light/dark cycle. Their accessory gland sizes not only exceeded the upper 95% CI limit of the newly eclosed males but also showed substantial growth in all strains (Figure 4A and Figure S1). Under the 8L:16D light/dark cycle, some individuals from the high- and mid-latitude strains went into reproductive arrest; however, the proportion did not exceed 60% in any strain, even under the condition where most of females from these regions entered reproductive diapause. These results suggest a weaker suppression of reproduction in males compared to females under unfavorable conditions. Consistent with the reproductive status of the females at 12°C, the males from low-latitude regions did not enter reproductive arrest under either photoperiod.

### Intraspecific variation in male genital morphology

To assess the degree of divergence among the latitudinal groups within this species, we examined another polymorphic trait related to reproduction that is not strongly influenced by climatic adaptation. Unlike the diapause-related traits, genital morphology in general is insensitive to environmental factors and is often used for species identification. Thus, variation in this trait provides an independent dataset for evaluating the extent of genetic divergence and independent segregation of genomic regions. Different morphological forms of aedeagus appendages have raised considerable debate over species delimitation of *D. triauraria*. Bock & Wheeler (1972) initially described *D. quadraria* from Taiwan as a distinct species based on differences in male genital morphology, specifically the presence of smaller appendages (dorsolateral claws) compared to those of *D. triauraria* (type specimen from Tokyo, Japan). However, subsequent fertility assays of hybrids (Kimura, 1987; Kim et al., 1989) and molecular phylogenetic analyses (Miyake & Watada, 2007) led to the reclassification of *D. quadraria* as a junior synonym of *D. triauraria* (Watada et al., 2011). Additionally, individuals with similarly small appendages have been reported from the southern islands of Japan (Onuma et al., 2022), further supporting that the observed variation reflects intraspecific rather than interspecific differences.

The morphological variation in aedeagus appendages was classified into two types: Northern (N) and Southern (S) (Figure 5 upper panel). Type N is characterized by larger appendages with a narrower space between them compared to type S. Type N corresponds to the originally described form in *D. triauraria* (Bock & Wheeler, 1972), while type S matches the form from Nansei (Ryukyu) Islands (Onuma et al., 2022), corresponding to the low-latitude region in this study. Genital morphology was examined in 10 males per strain. The strains from southern Japan (low-latitude, Yakushima, and two mid-latitude strains) possessed only type S, whereas most mid- and high-latitude strains included both types (Figure 5 lower panel). Although weak differentiation between northern and southern populations was apparent in genital morphology, discrepancies between the genital morphology and the diapause phenotype suggest that these traits segregate independently, consistent with long-term gene flow and mixed genomic backgrounds across geographic regions.

**FIGURE 5.**
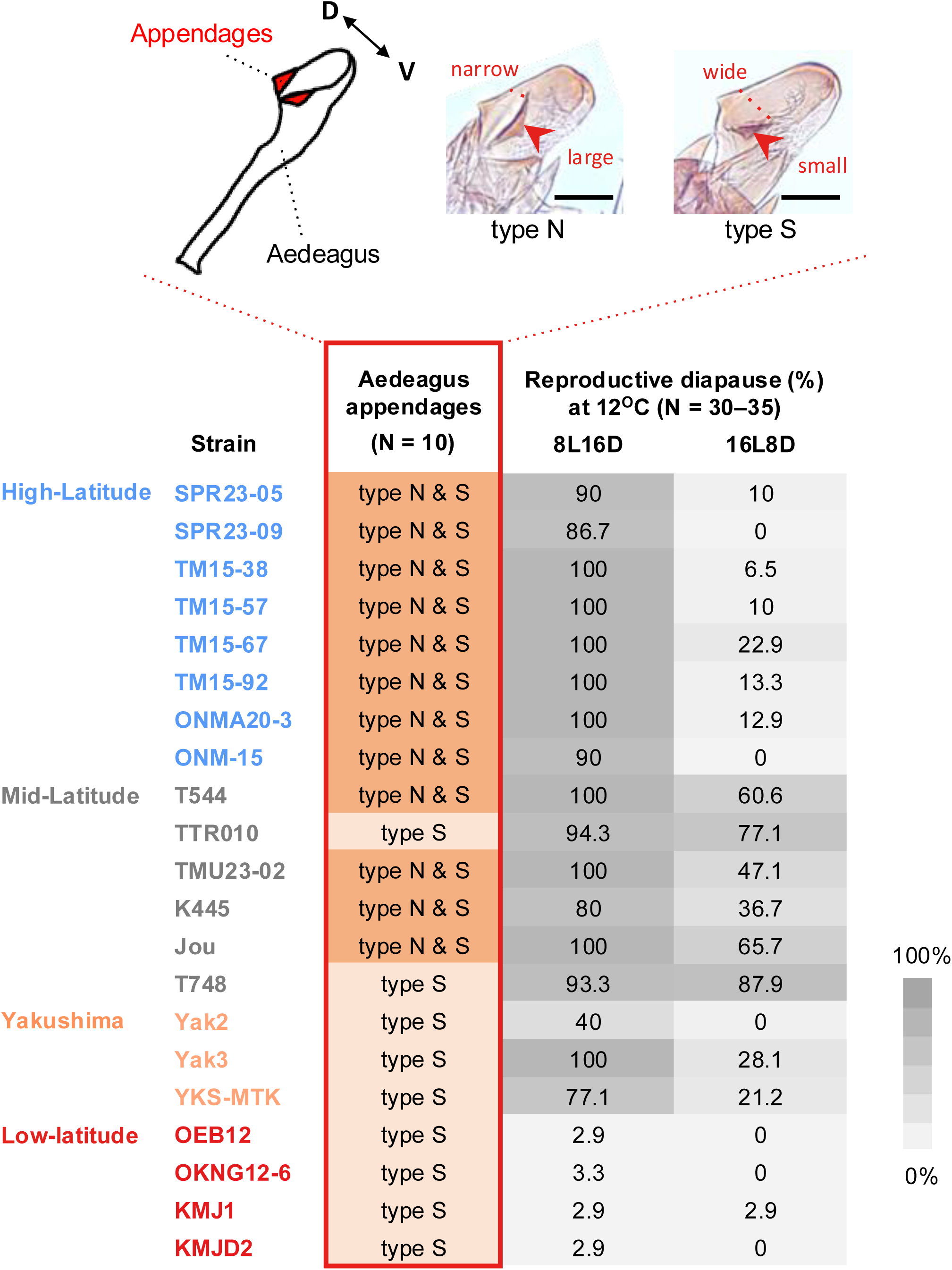
Polymorphism of the aedeagus appendages (dorsolateral claws) compared with female reproductive diapause phenotype in *D. triauraria*. The upper panel shows the schematic images of the size polymorphism of aedeagus appendages (after Onuma et al. 2022), classified as Northern (N) and Southern (S) types. Scale bar indicates 50 *μ*m. Genital morphology was examined in 10 males per strain and compared with the female reproductive diapause phenotype at 12°C (lower panel). The colors of the strain names represent their latitudinal regions of origin: blue, grey, orange, and red indicate high-latitude, mid-latitude, Yakushima, and low-latitude regions, respectively.

### Genomic phylogeny and geographic differentiation in *D. triauraria*

Whole-genome sequences were analyzed to further investigate the overall genetic differentiation among *D. triauraria* strains and the relationship to three other species from the auraria complex. Short-read sequencing data were generated from 14 *D. triauraria*, one *D. subauraria*, and one *D. biauraria* strains, and additional data for seven *D. triauraria* and four *D. auraria* strains were retrieved from the NCBI Sequence Read Archive (SRA) (Table 1). Phylogenetic analysis using a maximum-likelihood tree based on the whole-genome sequences revealed that strains from *D. triauraria* and *D. auraria* formed separate clades (Figure 6). Within *D. triauraria*, a weak north to south clinal differentiation was observed, with southern population being closer to the other three species of the auraria complex. The low-latitude strains formed a single cluster, whereas strains within each of the high-, mid-latitude, and Yakushima regions did not.

**FIGURE 6.**
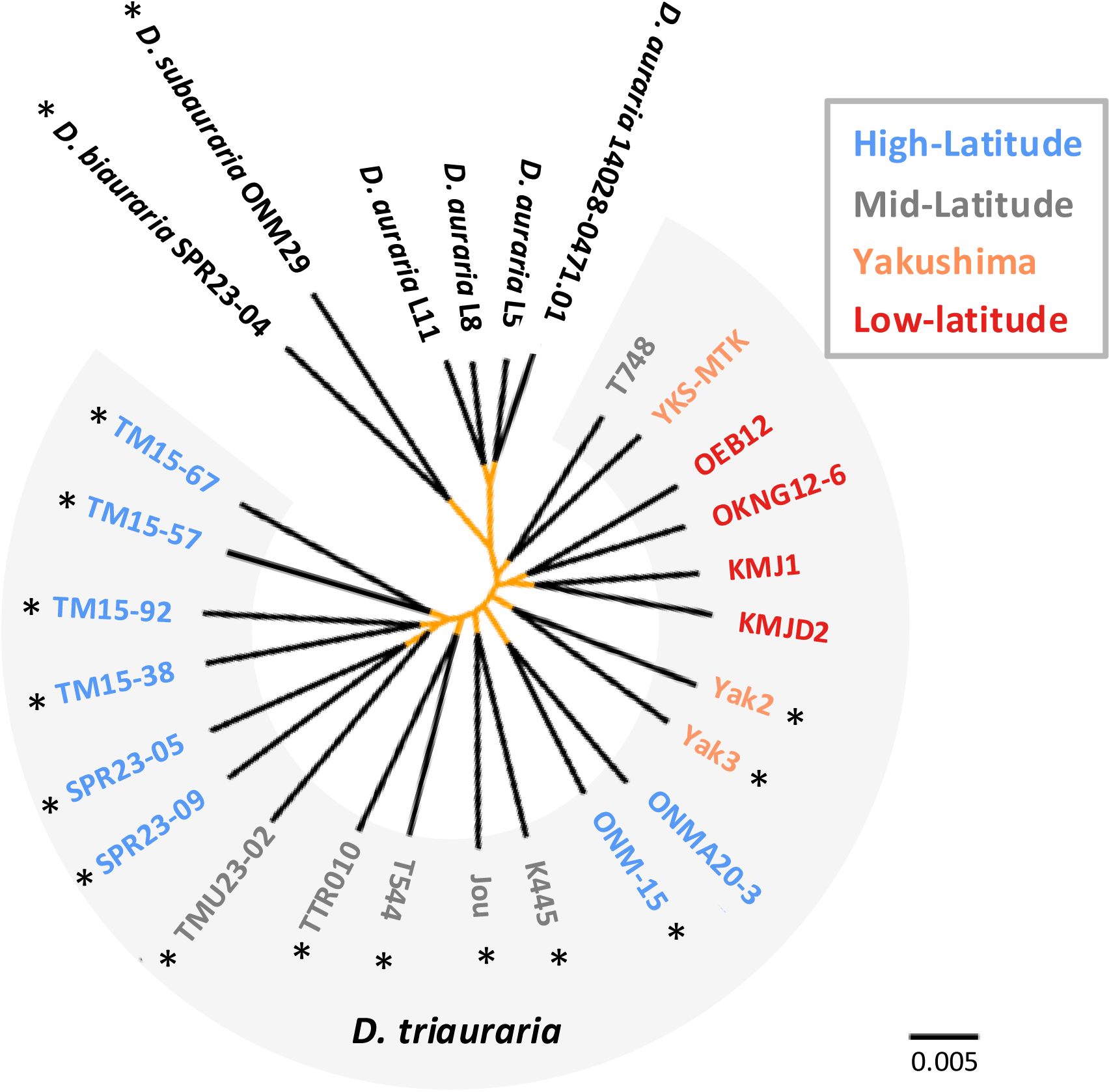
Maximum likelihood tree constructed from the whole-genome sequences of *D. triauraria, D. auraria, D. biauraria,* and *D. subauraria* strains used in this study. Maximum likelihood tree using IQ-TREE with the GTR+G+I model. Nodes with UFBoot value = 100, estimated from 1,000 pseudo-replicates are highlighted in orange. The colors of *D. triauraria* strain names represent their latitudinal regions of origin: blue, grey, orange, and red indicate high-latitude, mid-latitude, Yakushima, and low-latitude regions, respectively. Asterisks indicate strains whose genomic sequences were obtained in this study.

### Identification of genomic regions differentiated among geographic populations

The genital morphology polymorphism and the whole-genome phylogeny suggest that divergence among latitudinal groups at loci regulating reproductive diapause is likely to be maintained despite gene flow. Such loci should therefore exhibit signatures of divergence among latitudinal groups, in contrast to the rest of the genome. To identify these regions, we employed a tree-based method, focusing on strains from extremes of the species distribution (high- and low-latitude groups), where diapause phenotypes differ most strongly. The genome was partitioned into 5 kb windows with a 1 kb step size, and the windows were screened for those in which high- and low-latitude strains formed monophyletic clusters with ≥ 90% bootstrap (UFBoot) support. A window size of 5 kb was chosen after comparison with 10, 20, and 100 kb windows, which produced largely consistent UFBoot profiles (Figure 7 and Figure S2), indicating that the 5 kb window provided finer-scale resolution while retaining sufficient phylogenetic signal.

**FIGURE 7.**
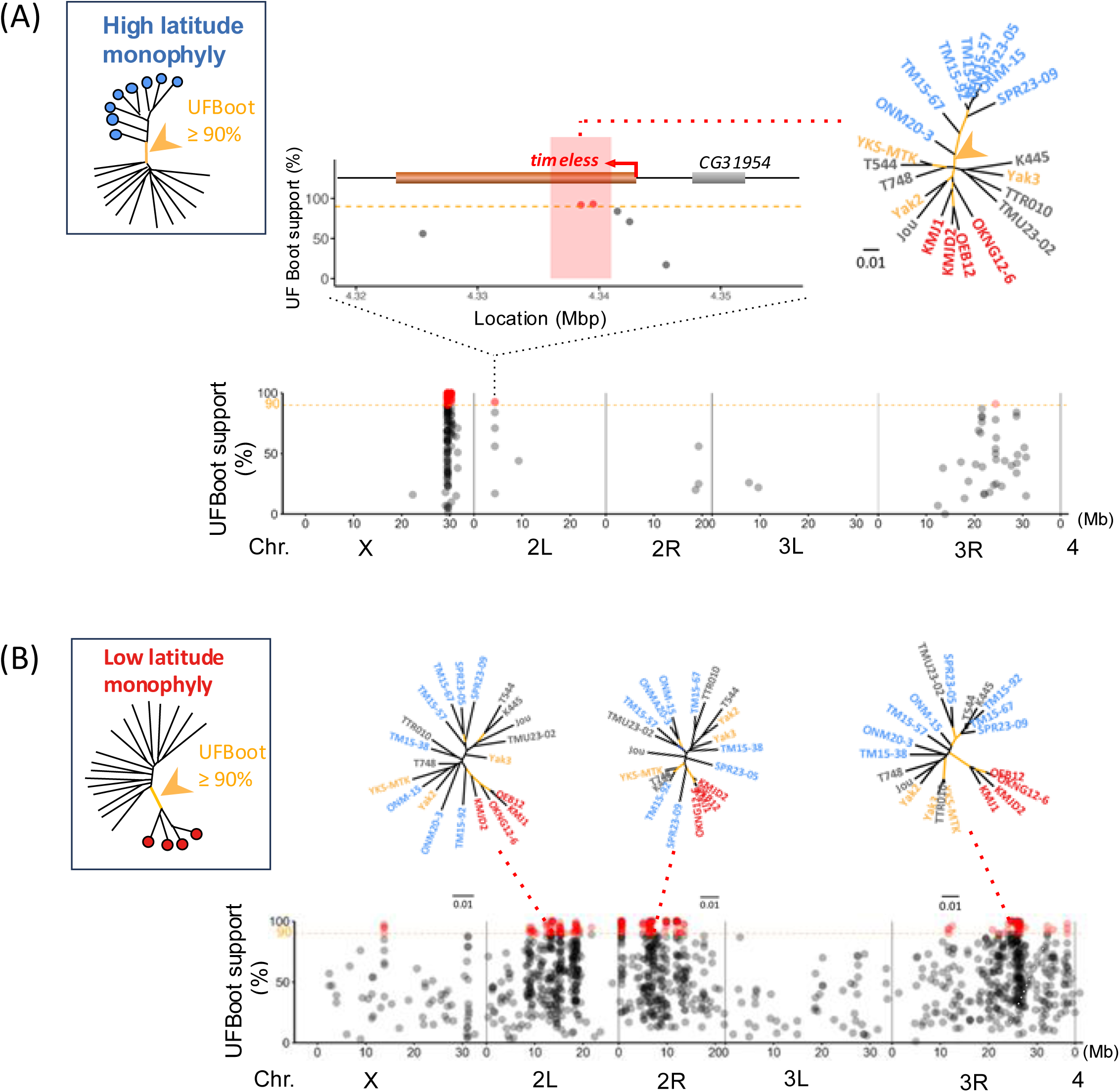
Monophyletic window analyses of the high- and low-latitudinal groups. (A) Sliding window profile of bootstrap (UFBoot) support (%) for monophyly of the high-latitude strain group, based on maximum parsimony trees (window size = 5 kb, step size = 1 kb). Red dots mark the midpoints of windows where the high-latitude strains formed monophyletic clusters with ≥ 90% bootstrap support (1,000 pseudo-replicates). The upper graph zooms in on the region containing the *timeless* (*tim*) gene (brown horizontal bar), which coincides with two consecutive windows above this threshold (midpoints in red). The maximum likelihood tree of one of these windows (highlighted by a pink stripe) is shown, with nodes having UFBoot ≥ 90% highlighted in orange. (B) Sliding-window profile of bootstrap (UFBoot) support (%) for monophyly of the low-latitude strain group, based on maximum parsimony trees (window size = 5 kb; step size = 1 kb). Red dots mark the midpoints of windows where low-latitude strains formed monophyletic clusters with ≥ 90% bootstrap support (1,000 pseudo-replicates). The maximum likelihood trees of the sampled windows are drawn above the graph, with nodes having UFBoot ≥ 90% highlighted in orange. The genome coordinates are based on the *D. triauraria* reference genome (GCA_014170255.2_RU_Dtri_1.1_genomic.fna). The colors of *D. triauraria* strain names represent their latitudinal origin: blue, grey, orange, and red indicate high-latitude, mid-latitude, Yakushima, and low-latitude regions, respectively.

For the high-latitude group, 55 monophyletic windows were detected on the X chromosome, two windows on 2L, and a single window on 3R (Figure 7A and Table S2). The clustered windows on the X chromosome, stretching discontinuously from 29.437 Mb to 30.287 Mb, included the region homologous to *D. melanogaster* Y chromosome (29.7–34.9 Mb, Chang et al., 2023) and therefore almost no coding genes were found (Table S2). Notably, the two windows on 2L included *timeless* (*tim*), which has been implicated for diapause regulation in many insect species (Mathias et al., 2005; Tauber et al., 2007; Sandrelli et al., 2007; Pruisscher et al., 2018; Lindestad et al., 2022; Vaze et al., 2024) and also suggested to be associated with diapause induction in *D. triauraria* (Yamada & Yamamoto, 2011). Sanger resequencing of the two consecutive windows spanning a 6 kb region confirmed the presence of two putative E-box motifs and one TER-box motif (Figure 8), which are critical for the circadian gene regulation (McDonald et al., 2001). The 11 polymorphic sites fixed in the high-latitude group were located in intronic regions outside these motifs.

**FIGURE 8.**
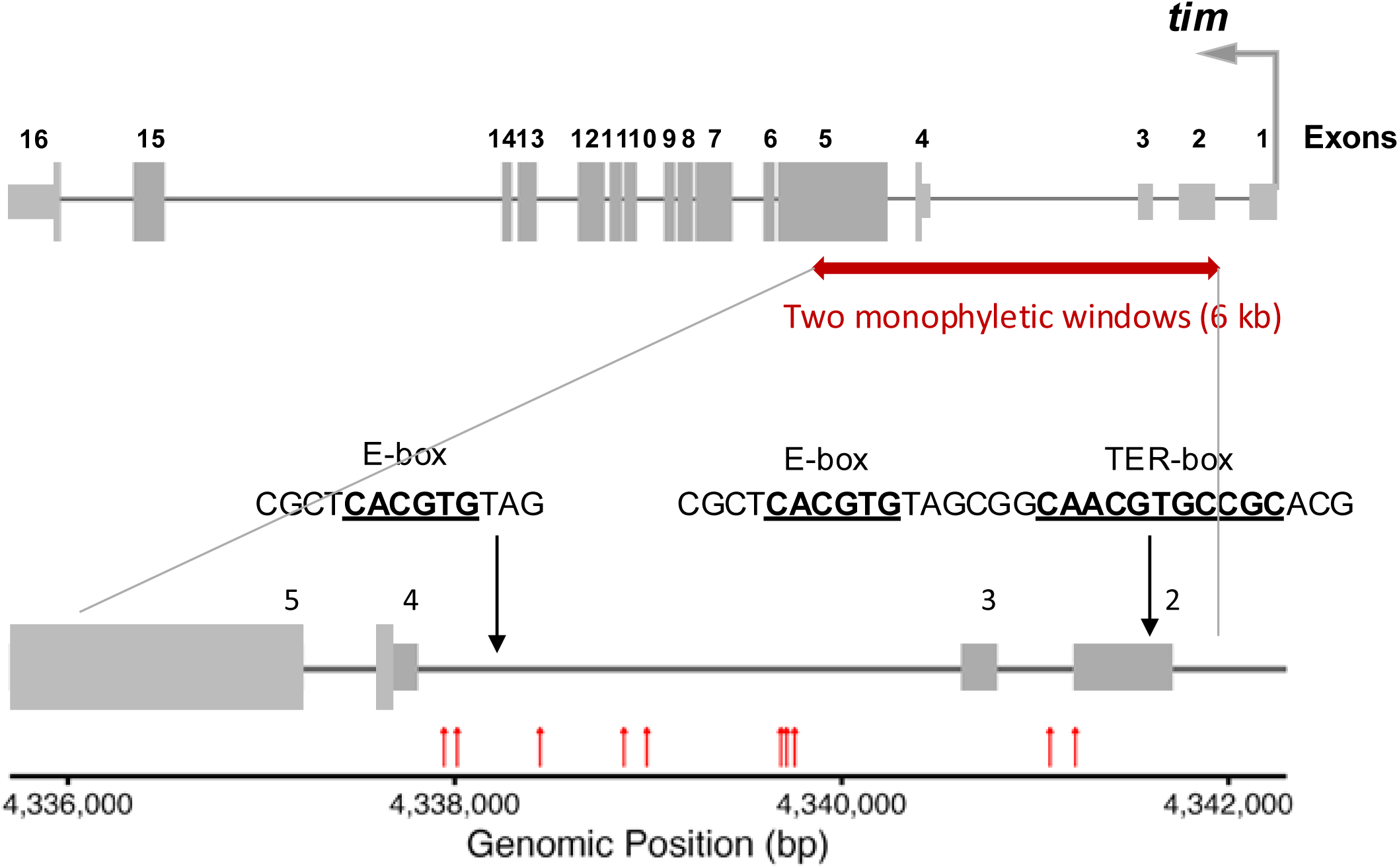
Putative E-box and TER-box sequence motifs within the 6 kb region covering the two consecutive monophyletic windows of the high-latitude strains. The two consecutive monophyletic 5 kb windows are shown in a red horizontal arrow. Gray boxes indicate predicted exons of *tim-L* isoform. Red vertical arrows mark the positions of SNPs that are fixed only in high-latitude populations (positions in bp: 4,337,946, 4,338,013, 4,338,442, 4,338,876, 4,338,994, 4,338,995, 4,339,687, 4,339,715, 4,339,759, 4,341,078, 4,341,209). Putative E-box and TER-box motifs are shown, with the positions corresponding to the indicated sites. The genome coordinates are based on the *D. triauraria* reference genome (GCA_014170255.2_RU_Dtri_1.1_genomic.fna).

In the low-latitude group, which does not respond to either photoperiod or temperature, a total of 188 monophyletic windows were located on chromosomes X, 2L, 2R, and 3R (Figure 7B, Table S3). These regions included *eyes shut* (*eys*), known to be involved in temperature entrainment of circadian activity (Sehadova et al., 2009); *nahoda*, which is expressed in egg chambers (Manning et al., 2017); and *Sox100B*; which plays a role in male-specific gonad development (Nanda et al., 2009). The involvement of these genes in the regulation of reproductive diapause has not been implicated.

The genome-wide F_ST_ window comparisons are more commonly applied to detect divergent genomic regions between populations. However, F_ST_ estimates can be biased when the sample sizes are small or uneven among regional populations (Whitlock and McCauley, 1999; Willing et al., 2012). To address this, we compared the windows with the highest F_ST_ values to those identified using the tree-based approach (Figure S3 and Figure S4). These comparisons revealed that 93.1% (54 out of 58) of the monophyletic windows in high-latitude strains (including those with the *tim* locus) and 92.0% (173 out of 188) in low-latitude strains overlapped with the top 5% F_ST_ windows. In contrast, only 0.09% (54 out of 5841) and 0.032% (173 out of 5463) of the top 5% windows overlapped with the monophyletic windows in high-and low-latitude strains, respectively. Thus, the monophyletic windows represent a stringently defined subset of the highest F_ST_ windows, based on an additional criterion of strong bootstrap support for monophyly.

### Association between the expression of *tim* isoform groups and diapause incidence

Multiple *tim* isoforms are expressed in *D. melanogaster* and other species of the *melanogaster* subgroup (Low et al., 2008; Anduaga et al., 2019). These isoforms can be categorized by size: long isoforms (*tim-L*, *tim-cold*), and a short isoform (*tim*-*sc*). *tim*-*L* is the canonical isoform, while *tim*-*cold* and *tim*-*sc* are more abundant at low temperatures. The roles of these isoforms have been investigated in *D. melanogaster*, variants of the *tim*-*L* isoform (*ls-tim*- and *s*-*tim*) are known to regulate diapause induction (Tauber et al., 2007; Sandrelli et al., 2007), and accumulation of TIM-SC under low temperature, short-day condition is implicated in diapause regulation by stabilizing Eyes absent (EYA) (Abrieux et al., 2020).

The isoform differences are mostly attributed to alternative splicing downstream of exon 9. Thus, to investigate the presence of these isoforms in *D. triauraria*, we performed 3′ RACE using RNA extracted from the heads of females from two strains, ONMA20-3 (high-latitude) and OEB12 (low-latitude). We were able to identify sequences corresponding to the two common isoforms, *tim*-*L* and *tim*-*sc*, and designed primers within the terminal exons to quantify the abundance of the long and short isoforms (hereafter referred to as *tim*-*L* and *tim*-*sc* isoform groups, respectively) by a realtime RT-qPCR (Figure S5). Although *tim*-*cold* and *tim*-*M* were not detected in our 3’ RACE analysis, they may still be amplified by the primers targeting the *tim*-*L* isoform group. It should be noted that other undetected isoforms may also be amplified by the designed primer pairs.

The oscillation patterns of *tim*-*L* and *tim*-*sc* isoform groups in the heads of females from the ONMA20-3 strain were analyzed under an 8L:16D or 16L:8D light/dark cycle at 12°C (Figure S6). The temporal patterns of *tim*-*L* expression under both photoperiods largely coincided with the nuclear accumulation of TIM in the lateral ventral neurons (LN_v_) of *D. melanogaster* brain (Shafer et al., 2004), including a possible phase advance at low temperature (Majercak et al., 1999; Anduaga et al., 2019). The overall expression level of *tim*-*L* isoform group (averaged across six timepoints) was significantly higher under the 16L:8D light/dark cycle than under 8L:16D (*t*-test after hyperbolic arcsine transformation: *t* = 14.94, df = 2, *P* < 0.01), whereas no significant difference was observed between the two cycles in the *tim*-*sc* isoform group (*t* = 3.54, df = 2, *P* = 0.07).

To investigate whether the differences between the two light conditions in expression of the two isoform groups (particularly *tim*-*L*, which showed a significant light-dependent difference in ONMA20-3) are associated with diapause incidence at 12°C, the *tim* mRNA abundance was quantified at ZT8 and ZT20 under 8L:16D and 16L:8D in female heads from the high-, mid-, and low-latitude strains (Figure S7A). Two-way ANOVA showed that the *tim*-*L* expression in the low-latitude strains was significantly lower than in the high-latitude strains under both light conditions and also significantly lower than in the mid-latitude strains under 8L:16D (Figure S7C), suggesting a distinct regulatory mechanism of this isoform group in the low-latitude strains. For the *tim*-*sc* isoform group, no significant effects were observed except for a difference between the high- and mid-latitude groups under the 8L:16D light/dark cycle (Figure S7B and Figure S7C).

Because the low-latitude strains exhibited no reproductive diapause even under the 8L:16D short-day (SD) condition at 12°C and displayed distinct *tim*-*L* expression profiles, the following association analysis was conducted using only the high- and mid-latitude strains, which showed variable diapause incidence under the 16L:8D long-day (LD) condition at 12°C. For simplicity, the abbreviations LD and SD are used in the model. To explain variation in diapause incidence under LD, we considered the following model:

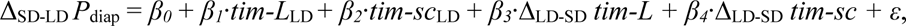

where Δ_SD-LD_ *P*_diap_ is the difference in the arcsine-transformed diapause proportions between the SD (8L:16D) and LD (16L:8D) conditions. *tim-L*_LD_ and *tim-sc*_LD_ denote the hyperbolic arcsine-transformed relative expression levels under LD, and Δ_LD-SD_ *tim-L* and Δ_LD-SD_ *tim-sc* represent the differences in expression between the two light conditions. The arcsine transformation was applied to normalize the diapause incidence, which ranges from 0 to 1, while the hyperbolic arcsine transformation was used to normalize the *tim* expression data, which are not restricted to this range. *β_0_* and *β_1_*_–*4*_ denotes intercept and regression coefficients, respectively. This model aims to explain the variable Δ_SD-LD_ *P*_diap_, reflecting the increased reproduction (reduced diapause) under LD relative to SD, which is most pronounced in the high-latitude strains (Figure 3). It incorporates *tim-L* and *tim-sc* isoform group expression under LD (*tim-L*_LD_ and *tim-sc*_LD_), as well as their expression differences between LD and SD (Δ_LD-SD_ *tim-L* and *β_4_*·Δ_LD-SD_ *tim-sc*), to account for the relationship between isoform-specific expression and photoperiodic reproductive response.

We then performed stepwise model selection using BIC, which identified the following model as the best-fit model:

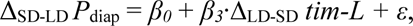

with the coefficient estimates shown in Table 2. The result indicates that the differences in *tim*-*L* expression between the two light conditions are significantly associated with the differences in diapause incidence at 12°C. The positive regression coefficient indicates that strains with relatively higher *tim-L* expression under the long-day condition (LD: 16L:8D) show reduced diapause incidence, suggesting an enhanced reproductive activity at low temperature.

**Table 2.**
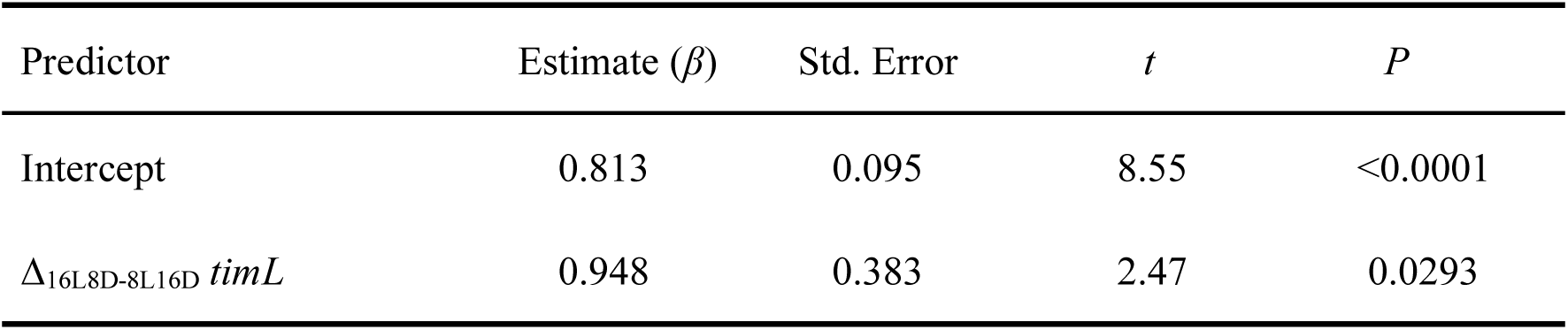
Results of the linear regression model selected by stepwise BIC for explaining the differences in diapause incidence between light conditions.

| Predictor | Estimate ($\beta$) | Std. Error | $t$ | $P$ |
| --- | --- | --- | --- | --- |
| Intercept | 0.813 | 0.095 | 8.55 | <0.0001 |
| $\Delta_{16L8D-8L16D}$ *timL* | 0.948 | 0.383 | 2.47 | 0.0293 |

## DISCUSSION

Initiating or arresting reproduction is a critical life-history choice with major consequences for lifetime fitness. Like many other insects, *D. triauraria* respond to environmental cues such as temperature and photoperiod to make the choice. We conducted a comprehensive analysis of the female reproductive diapause induction in 21 strains of this species, including both newly tested and previously reported strains collected from across the Japanese archipelago (26.2–43.1°N). Our results revealed that a key difference between the high- and mid-latitude strains lies in their temperature sensitivity under the long-day condition (16L:8D). At 12°C, females from the mid-latitude strains exhibited a higher incidence of reproductive diapause compared to those from the high-latitude strains (Figure 3). This reduced sensitivity to low temperature (i.e., stronger photoperiodic induction of diapause) in the high-latitude strains likely reflects adaptation to northern environments, where reproducing under cooler and shorter summer seasons is essential.

Although direct comparisons between sexes are challenging, our results suggest that females and males from the high- and mid-latitude regions differ in how reproduction activity is modulated by photoperiod and temperature (Figure 3 and Figure 4). This implies that the thresholds for reproductive suppression vary between sexes, potentially reflecting sex-specific life history strategies. In particular, males may maintain investment in reproduction under suboptimal conditions, consistent with the relatively lower energetic cost of reproduction compared to females (Bateman, 1948; Trivers, 1972). Future work should examine the extent to which shared versus sex-specific neuroendocrine pathways and sensory circuits contribute to these differences. Moreover, assessing traits beyond accessory gland development, such as spermatogenesis status and courtship behavior, will be essential for a comprehensive understanding of intersexual variation in reproductive status under unfavorable environmental conditions.

Our whole-genome phylogeny revealed only weak geographic divergence (Figure 6), and the observed discrepancies between genital morphology and diapause phenotype (Figure 5) further suggest that long-term gene flow across geographic regions has generated sufficient recombination to decouple genomic regions associated with different traits. Such genomic decoupling provides an opportunity to identify geographically divergent regions associated with specific phenotypes. To investigate the genetic basis underlying the observed differences in female diapause response, we applied a monophyletic window approach, which imposes a stricter criterion for identifying genomic regions than the standard F_ST_ scans. Although Weir and Cockerham’s weighted F_ST_ estimation corrects for sample sizes of the populations (Weir and Cockerham 1984), small and unbalanced samples, as in this study, can introduce false signals of differentiation (Whitlock and McCauley, 1999; Willing et al., 2012). By focusing on genomic windows where the focal group forms a well-supported monophyletic cluster, this approach helps mitigate such biases.

For the high-latitude strain group (against all other groups), only 54 5-kb windows showed both ≥ 90% bootstrap support for monophyly and F_ST_ values within the top 5%, representing <1% of all top 5% F_ST_ windows (Figure S4A). These windows were located in two separate genomic regions (one additional window on 3R met the monophyly criterion but fell outside the top 5% F_ST_ threshold) (Figure 7A). The two windows on 2L contained *tim*, and a cluster of windows on the X chromosome included few genes (Table S1). For the low-latitude strain group, 173 windows met both the ≥ 90% bootstrap support and top 5% F_ST_ criteria, representing approximately 3% of all top 5% F_ST_ windows (Figure S4B). These were distributed across more than 20 separate genomic regions (Figure 7B). The smaller number of candidate windows in the low-latitude groups could be due to multiple factors: a smaller number of strains, a higher number of regions under selection, inflated F_ST_ threshold in the high-latitude strains due to a dense cluster of high-F_ST_ windows on the X chromosome, or a combination of these factors. We were unable to examine specific candidate regions within the low-latitude group due to these complexities.

Our monophyletic window approach does not directly target diapause-associated loci and may capture divergent regions associated with other locally adapted traits, including non-reproductive traits such as the recently reported midgut remodeling response during diapause in this species (Adachi et al. 2025), as well as more general metabolic or stress-tolerance adaptations. Nevertheless, among the windows showing strong signals of high-latitude group monophyly, we found a candidate region encompassing the E-box and T-box regulatory motifs of *tim*. Given the implicated role of *tim* in regulating diapause across multiple insect species, including *D. triauraria*, we further investigated the association between *tim* expression levels and reproductive diapause incidence. Using RT-qPCR, the mRNA levels of the long and short isoform groups were quantified. We detected a significant association across strains between the differences in diapause incidence under the long- and short-day photoperiods and the differences of *tim-L* isoform group expression between those light conditions at 12°C. The analysis focused on the high- and mid-latitude strains, as the low-latitude strain exhibited distinct diapause regulation patterns. Finally, it would be informative in future studies to quantify other non-reproductive stress-response traits under these conditions, as they may also be regulated by *tim*.

As in many organisms, circadian rhythms in *Drosophila* are influenced by light, implicating a link between clock genes and photoperiodic regulation (e.g., Denlinger, 2022). *tim* is a core circadian gene whose product, TIM, is bound by the photoreceptor CRYPTOCHROME (CRY) upon light exposure and subsequently degraded by ubiquitination, resulting in light-induced resetting of the circadian clock (Hunter-Ensor et al., 1996; Ceriani et al., 1999; Busza et al., 2004). The primary light-sensitive isoform, *tim*-*L*, is included in the isoform group for which we observed an association with diapause incidence. The light-insensitive isoforms, *tim*-*cold* and *tim*-*sc*, are induced by low temperature (Montelli et al., 2015; Shakhmantsir et al., 2018; Anduaga et al., 2019; Abrieux et al., 2020). Among these isoforms, TIM-SC contributes to the stabilization of EYA, a key seasonal sensor, suggesting that the relative abundances of *tim*-*L* and *tim*-*sc* may play a role in diapause induction in females of *D. melanogaster* (Abrieux et al., 2020). There is no evidence from these *D. melanogaster* studies, however, demonstrating a direct effect of *tim*-*L* on reproductive diapause. Thus, the mechanisms underlying the observed association in our study, where higher *tim*-*L* expression associates with lower diapause incidence at low temperature, remains unclear at this point. It is also important to note that the TIM protein abundance was not quantified in our study, and the increased mRNA expression observed under the long-day condition could reflect either an enhanced transcription or compensatory response to light-driven protein degradation. Interestingly, unlike *D. melanogaster* and related species such as *D. simulans, D. yakuba,* and *D. virilis*, whose *tim* splicing is temperature-sensitive (Anduaga et al., 2019), the cold-tolerant northern species, *D. montana*, exhibits photoperiod-sensitive *tim* splicing (Tapanainen et al., 2018). This suggests that the splicing regulation of *tim* isoforms and their functional roles in circadian and diapause pathways may vary considerably across species.

In *D. triauraria*, Kimura and Yoshida (1995) analyzed recombinant inbred lines derived from hybrids between a diapausing strain (ONMA20-3) and a non-diapausing strain from Kametoku (28°N), and estimated that three or four loci underly the difference in the photoperiodic response, with at least one located on the X chromosome. More recently, Yamada and Yamamoto (2011) conducted a genetic analysis and reported an association between reproductive diapause incidence and the genetic marker variants at *tim* and *cry*. This analysis was based on genotyping five clock genes (*per*, *tim*, *clk*, *cyc*, and *cry*) in 329 backcrossed individuals derived from a cross between a diapausing strain (ONMA20-3) and a non-diapausing strain (OEB12). The association with *tim* is consistent with our results, further supporting a potential role for this gene in diapause regulation of *D. triauraria*. However, the low marker density (only five loci across the entire genome) limits their mapping resolution. As they also noted, the observed associations could be due to linkage with nearby genes rather than direct effects of *tim* or *cry*. Our higher-resolution approach allows a more targeted investigation of this region and together with the expression analysis, strengthens the case for *tim* as a candidate gene involved in photoperiodic diapause regulation.

Consistent with the previous report (Yamada & Yamamoto, 2011), females of low-latitude strains did not exhibit reproductive diapause under any of the tested conditions (Figure 3). Our experimental design differed from that of Yamada and Yamamoto (2011) in that flies were transferred to the test conditions immediately after eclosion, whereas in the earlier study flies were exposed to experimental conditions from the embryonic stage. Despite this difference, we observed qualitatively similar phenotypes in the low-latitude strains. We further showed that males of these strains also failed to arrest reproduction (Figure 4) and displayed reduced expression of *tim*-*L* isoform group (Figure S7A). Taken together with the earlier finding that the diapausing phenotype is completely dominant over the non-diapausing type (Yamada & Yamamoto, 2011), these results suggest that the low-latitude strains may carry impaired signaling pathways required for diapause induction. An intriguing evolutionary question that remains unresolved is the ancestry of the phenotype; whether the absence of diapause in the low-latitude strains represents a derived loss of the trait, or alternatively, whether the ability to enter reproductive diapause was an adaptation acquired in the high-latitude populations.

Many *Drosophila* species are likely to have originated in tropical region and later expanded into temperate and subarctic regions (Throckmorton, 1975; Lemeunier et al., 1986; Markow & O’Grady, 2006). In our whole-genome maximum likelihood tree, the low-latitude and Yakushima strains of *D. triauraria* form the basal lineages (Figure 6), consistent with a tropical origin. However, Kimura (1988) argued that adaptation to cool-temperate climates likely preceded speciation within the *auraria* complex. This view was based on the observation that species within the complex do not differ markedly in cold-hardiness. Supporting this hypothesis, three other species in the complex exhibit reproductive diapause phenotypes similar to that of the northern strains of *D. triauraria* (Kimura, 1984; Kimura, 1990). Kimura (1988) further proposed that the current subtropical populations of *D. triauraria* may have established through reinvasion from northern, cool-temperate populations, as these subtropical flies are significantly more cold-hardy than sympatric populations of the subtropical or warm-temperate species like *D. rufa* and *D. lutescens*. These competing hypotheses, tropical origin with independent acquisition of diapause in the north versus ancestral diapause with secondary loss in the south, remain difficult to reconcile.

However, the phylogenetic analysis of the *tim* genomic region, including other species of the auraria complex (Figure S8), does not support a scenario in which the low- and mid-latitude alleles are derived from the high-latitude allele, nor the reverse. Instead, the tree suggests that these alleles have diverged independently from a basal state. Given the likelihood of frequent introgression and incomplete lineage sorting within the *auraria* complex (Miyake & Watada, 2007; Conner et al., 2021), it is plausible that two alleles of early origin associated with diapause regulation have persisted in different geographic regions within *D. triauraria*. Although our analysis is currently confined to a single locus, the results raise the possibility that key diapause-related alleles originated and diverged close to the time of speciation and subsequently underwent differentiation along latitudinal gradients. In light of the polygenic basis of diapause regulation, information from additional loci will be essential to further test and refine this scenario, thereby providing deeper insights into the evolutionary dynamics of adaptive traits evolving across heterogeneous environments.

## ACKNOWLEDGEMENTS

We thank KYORIN-Fly stock center, KYOTO *Drosophila* stock center, and Shin G. Goto for providing *D. triauraria* stocks, and Masahito T. Kimura for providing *D. triauraria* and *D. biauraria* flies. We are grateful to Koichiro Tamura for fruitful discussions and support. This work was supported by KAKENHI 23K27221 to AT, 24KJ0181 to MO, 22H05073 to MN, and 22KJ2552 to TF. We used ChatGPT-4.0 to assist in correcting errors in script writing.

## AUTHOR CONTRIBUTIONS

TF and AT conceived this work. YA and TF analyzed accessory gland. MO and TF analyzed genital morphology. MN obtained genome sequences of five *D. triauraria* strains. TF and AT analyzed data and wrote the manuscript.

## DATA AVAILABILITY STATEMENT

Raw sequence reads have been deposited in the SRA under BioProject PRJDB20728, with individual run accession numbers provided in Table 1. Sequences of *tim* isoform groups are available in DDBJ (accession numbers LC882409–LC882412). Sequencing data for the 6 kb *tim* region are deposited in DDBJ (accession numbers LC884490–LC884507)

## REFERENCES

Abrieux, A., Xue, Y., Cai, Y., Lewald, K. M., Nhu Nguyen, H., Zhang, Y., Chiu, J. C. (2020). EYES ABSENT and TIMELESS integrate photoperiodic and temperature cues to regulate seasonal physiology in *Drosophila*. Proceedings of the National Academy of Sciences of the United States of America, 117(26), 15293–15304. 10.1073/pnas.2004262117

Adachi, Y., Nagai, H., Fujichika, T., Takahashi, A., Miura, M., & Nakajima, Y. (2025). BubR1 and Mad2 regulate adult midgut remodeling in *Drosophila* diapause [Preprint]. bioRxiv. 10.1101/2025.01.26.634899

Ala-Honkola, O., Kauranen, H., Tyukmaeva, V., Boetzl, F. A., Hoikkala, A., & Schmitt, T. (2020). Diapause affects cuticular hydrocarbon composition and mating behavior of both sexes in *Drosophila montana*. Insect Science, 27(2), 304–316. 10.1111/1744-7917.12639

Alexander, D. H., Novembre, J., & Lange, K. (2009). Fast model-based estimation of ancestry in unrelated individuals. Genome Research, 19(9), 1655–1664. 10.1101/gr.094052.109

Altschul, S. F., Gish, W., Miller, W., Myers, E. W., & Lipman, D. J. (1990). Basic local alignment search tool. Journal of Molecular Biology, 215(3), 403–410. 10.1016/S0022-2836(05)80360-2

Andrews, S. (2010). FastQC: A quality control tool for high throughput sequence data [Online]. Available at: http://www.bioinformatics.babraham.ac.uk/projects/fastqc/

Anduaga, A. M., Evanta, N., Patop, I. L., Bartok, O., Weiss, R., & Kadener, S. (2019). Thermosensitive alternative splicing senses and mediates temperature adaptation in *Drosophila*. eLife, 8, e44642. 10.7554/eLife.44642

Barghi, N., Hermisson, J., & Schlötterer, C. (2020). Polygenic adaptation: a unifying framework to understand positive selection. Nature Reviews Genetics, 21(12), 769–781. 10.1038/s41576-020-0250-z

Bateman, A. (1948). Intra-sexual selection in *Drosophila*. Heredity, 2, 349–368 10.1038/hdy.1948.21

Bock, I. R., & Wheeler, M. R. (1972). The *Drosophila melanogaster* species group. University of Texas Publications.

Boom, R., Sol, C. J. A., Salimans, M. M., Jansen, C. L., Wertheim-van Dillen P. M. E., & van der Noordaa J. (1990). Rapid and simple method for purification of nucleic acids. Journal of Clinical Microbiology, 28(3) 495–503. 10.1128/jcm.28.3.495-503.1990

Bradshaw, W. E., Emerson, K. J., Catchen, J. M., Cresko, W. A., & Holzapfel, C. M. (2012). Footprints in time: comparative quantitative trait loci mapping of the pitcher-plant mosquito, *Wyeomyia smithii*. Proceedings of the Royal Society B: Biological Sciences, 279(1747), 4551–4558. 10.1098/rspb.2012.1917

Bradshaw, W. E., & Holzapfel, C. M. (2007). Evolution of animal photoperiodism. Annual Review of Ecology, Evolution, and Systematics, 38, 1–25. 10.1146/annurev.ecolsys.37.091305.110115

Busza, A., Emery-Le, M., Rosbash, M., & Emery, P. (2004). Roles of the two *Drosophila* CRYPTOCHROME structural domains in circadian photoreception. Science, 304(5676), 1503–1506. 10.1126/science.1096973

Camacho, C., Coulouris, G., Avagyan, V., Ma, N., Papadopoulos, J., Bealer, K., & Madden, T. L. (2009). BLAST+: architecture and applications. BMC Bioinformatics, 10, 421. 10.1186/1471-2105-10-421

Ceriani, M. F., Darlington, T. K., Staknis, D., Más, P., Petti, A. A., Weitz, C. J., & Kay, S. A. (1999). Light-dependent sequestration of TIMELESS by CRYPTOCHROME. Science, 285(5427), 553–556. 10.1126/science.285.5427.553

Chang, C. H., Natividad, I. M., & Malik, H. S. (2023). Expansion and loss of sperm nuclear basic protein genes in *Drosophila* correspond with genetic conflicts between sex chromosomes. eLife, 12:e85249. 10.7554/eLife.85249

Chang, V., & Meuti, M. E. (2020). Circadian transcription factors differentially regulate features of the adult overwintering diapause in the Northern house mosquito, *Culex pipiens*. Insect Biochemistry and Molecular Biology, 121, 103365. 10.1016/j.ibmb.2020.103365

Conner, W. R., Delaney, E. K., Bronski, M. J., Ginsberg, P. S., Wheeler, T. B., Richardson, K. M., Peckenpaugh, B., Kim, K. J., Watada, M., Hoffmann, A. A., Eisen, M. B., Kopp, A., Cooper, B. S. & Turelli, M. (2021). A phylogeny for the *Drosophila montium* species group: A model clade for comparative analyses. Molecular Phylogenetics and Evolution, 158, 107061. 10.1016/j.ympev.2020.107061

Danecek, P., Auton, A., Abecasis, G., Albers, C. A., Banks, E., DePristo, M. A., Handsaker, R. E., Lunter, G., Marth, G. T., Sherry, S. T., McVean, G., Durbin, R., & 1000 Genomes Project Analysis Group. (2011). The variant call format and VCFtools. Bioinformatics, 27(15), 2156–2158. 10.1093/bioinformatics/btr330

Danecek P., Bonfield J. K., Liddle J., Marshall J., Ohan V., Pollard M. O., Whitwham A., Keane T., McCarthy S. A., Davies R. M., & Li H. (2021). Twelve years of SAMtools and BCFtools. GigaScience, 10(2), giab008. 10.1093/gigascience/giab008

Denlinger, D. L. (2022). Insect diapause. Cambridge University Press.

Denlinger, D. L. (2023). Insect diapause: from a rich history to an exciting future. Journal of Experimental Biology, 226(4): jeb245329. 10.1242/jeb.245329

Drummond-Barbosa, D., & Spradling, A. C. (2001). Stem cells and their progeny respond to nutritional changes during *Drosophila* oogenesis. Developmental Biology, 231(1), 265– 278. 10.1006/dbio.2000.0135

Emerson, K. J., Dake, S. J., Bradshaw, W. E., & Holzapfel, C. M. (2009). Evolution of photoperiodic time measurement is independent of the circadian clock in the pitcher-plant mosquito, *Wyeomyia smithii*. Journal of Comparative Physiology A, 195(4), 385– 391. 10.1007/s00359-009-0416-9

Ewels, P., Magnusson, M., Lundin, S., & Käller, M. (2016). MultiQC: summarize analysis results for multiple tools and samples in a single report. Bioinformatics, 32(19), 3047–3048. 10.1093/bioinformatics/btw354

Giorgi, F., & Deri, P. (1976). Cell death in ovarian chambers of *Drosophila melanogaster*. Journal of Embryology and Experimental Morphology, 35(3), 521–533. 10.1242/dev.35.3.521

Haas, B. J., Papanicolaou, A., Yassour, M., Grabherr, M., Blood, P. D., Bowden, J., Couger, M. B., Eccles, D., Li, B., Lieber, M., Macmanes, M. D., Ott, M., Orvis, J., Pochet, N., Strozzi, F., Weeks, N., Westerman, R., William, T., Dewey, C. N., Henschel, R., LeDuc, R. D., Friedman, N., & Regev, A. (2013). De novo transcript sequence reconstruction from RNA-seq using the Trinity platform for reference generation and analysis. Nature Protocols, 8(8), 1494–1512. 10.1038/nprot.2013.084

Hall, T. A. (1999). BioEdit: a user-friendly biological sequence alignment editor and analysis program for Windows 95/98/NT. Nucleic Acids Symposium Series, 41, 95–98.

Hasebe, M., & Shiga, S. (2025). A Peptidergic Neural system connects the circadian clock to the photoperiodic control of reproductive diapause in the bug *Riptortus pedestris*. The Journal of Neuroscience, 45(50), e1717252025. 10.1523/JNEUROSCI.1717-25.2025

Hoang, D. T., Vinh, L. S., Flouri, T., Stamatakis, A., von Haeseler, A., & Minh, B. Q. (2018). MPBoot: Fast phylogenetic maximum parsimony tree inference and bootstrap approximation. BMC Evolutionary Biology, 18(1), 11. 10.1186/s12862-018-1131-3

Hunter-Ensor, M., Ousley, A., & Sehgal, A. (1996). Regulation of the *Drosophila* protein timeless suggests a mechanism for resetting the circadian clock by light. Cell, 84(5), 677–685. 10.1016/S0092-8674(00)81046-6

Hut, R. A., Paolucci, S., Dor, R., Kyriacou, C. P., & Daan, S. (2013). Latitudinal clines: an evolutionary view on biological rhythms. Proceedings of the Royal Society B: Biological Sciences, 280(1765), 20130433. 10.1098/rspb.2013.0433

Ichijo, N. (1986). Disjunctive cline of critical photoperiod in the reproductive diapause of *Drosophila lacertosa*. Evolution, 40(2), 418–421. 10.2307/2408820

Ikeno, T., Tanaka, S. I., Numata, H., & Goto, S. G. (2010). Photoperiodic diapause under the control of circadian clock genes in an insect. BMC biology, 8, 116. 10.1186/1741-7007-8-116

Johnson, O. L., Tobler, R., Schmidt, J. M., & Huber, C. D. (2023). Fluctuating selection and the determinants of genetic variation. Trends in Genetics, 39(6), 491–504. 10.1016/j.tig.2023.02.004

Kim, B. K., Watanabe, T. K., & Kitagawa, O. (1989). Evolutionary genetics of the *Drosophila montium* subgroup. I. Reproductive isolations and the phylogeny. Japanese Journal of Genetics, 64(3), 177–190. 10.1266/jjg.64.177

Kim, D., Paggi, J. M., Park, C., Bennett, C., & Salzberg, S. L. (2019). Graph-based genome alignment and genotyping with HISAT2 and HISAT-genotype. Nature Biotechnology, 37(8), 907–915. 10.1038/s41587-019-0201-4

Kimura, M. T. (1983). Geographic variation and genetic aspects of reproductive diapause in *Drosophila triauraria* and *D. quadraria*. Physiological Entomology, 8(2), 181–186. 10.1111/j.1365-3032.1983.tb00347.x

Kimura, M. T. (1984). Geographic variation of reproductive diapause in the *Drosophila auraria* complex (Diptera: Drosophilidae). Physiological Entomology, 9(4), 425–431. 10.1111/j.1365-3032.1984.tb00784.x

Kimura, M. T. (1987). Habitat differentiation and speciation in the *Drosophila auraria* species-complex (Diptera, Drosophilidae). Kontyu, 55(3), 429–436.

Kimura, M. T. (1988a). Adaptations to temperate climates and evolution of overwintering strategies in the *Drosophila melanogaster* species group. Evolution, 42(6), 1288–1297. 10.1111/j.1558-5646.1988.tb04188.x

Kimura, M. T. (1988b). Interspecific and geographic variation of diapause Intensity and seasonal adaptation in the *Drosophila auraria* species complex (Diptera: Drosophilidae). Functional Ecology, 2(2), 177–183. 10.2307/2389693

Kimura, M. T. (1990). Quantitative response to photoperiod during reproductive diapause in the *Drosophila auraria* species-complex. Journal of Insect Physiology, 36(3), 147–152. 10.1016/0022-1910(90)90115-V

Kimura, M. T., & Yoshida, T. (1995). A genetic analysis of photoperiodic reproductive diapause in *Drosophila triauraria*. Physiological Entomology, 20(3), 253–256. 10.1111/j.1365-3032.1995.tb00009.x

King, R. C. (1970). Ovarian development in *Drosophila melanogaster*. Academic Press

Krueger, F. (2015). Trim Galore! [Online]. Available at: https://www.bioinformatics.babraham.ac.uk/projects/trim_galore/

Kozak, G. M., Wadsworth, C. B., Kahne, S. C., Bogdanowicz, S. M., Harrison, R. G., Coates, B. S., & Dopman, E. B. (2019). Genomic basis of circannual rhythm in the European corn borer moth. Current Biology, 29(20), 3501–3509.e5. 10.1016/j.cub.2019.08.053

Kubrak, O. I., Kucerová, L., Theopold, U., Nylin, S., & Nässel, D. R. (2016). Characterization of reproductive dormancy in male *Drosophila melanogaster*. Frontiers in Physiology, 7, 572. 10.3389/fphys.2016.00572

Kumar, S., Stecher, G., & Tamura, K. (2016). MEGA7: Molecular evolutionary genetics analysis version 7.0 for bigger datasets. Molecular Biology and Evolution, 33(7), 1870–1874. 10.1093/molbev/msw054

Lankinen, P. (1986). Geographical variation in circadian eclosion rhythm and photoperiodic adult diapause in *Drosophila littorlis*. Journal of comparative physiology A, 159, 123– 142. 10.1007/BF00612503

Lankinen, P., Kastally, C., & Hoikkala, A. (2023). Clinal variation in the temperature and photoperiodic control of reproductive diapause in *Drosophila montana* females. Journal of Insect Physiology, 150, 104556. 10.1016/j.jinsphys.2023.104556

Langmead, B., & Salzberg, S. L. (2012). Fast gapped-read alignment with Bowtie 2. Nature Methods, 9(4), 357–359. 10.1038/nmeth.1923

Láruson, Á. J., Yeaman, S., & Lotterhos, K. E. (2020). The importance of genetic redundancy in evolution. Trends in Ecology and Evolution, 35(9), 809–822. 10.1016/j.tree.2020.04.009

Lee, S. F., Sgro, C. M., Shirriffs, J., Wee, C. W., Rako, L., Heerwaarden, B. V., & Hoffmann, A. A. (2011). Polymorphism in the *couch potato* gene clines in eastern Australia but is not associated with ovarian dormancy in *Drosophila melanogaster*. Molecular Ecology, 20(14), 2973–2984. 10.1111/j.1365-294X.2011.05155.x

Lemeunier, F., David, J., Tsacas, L., & Ashburner, M. (1986). The *melanogaster* species group. genetics and biology of *Drosophila* 3e, Academic Press.

Li, H., Handsaker, B., Wysoker, A., Fennell, T., Ruan, J., Homer, N., Marth, G., Abecasis, G., & Durbin, R. (2009). The sequence alignment/map format and SAMtools. Bioinformatics, 25(16), 2078–2079. 10.1093/bioinformatics/btp352

Lindestad, O., Nylin, S., Wheat, C. W., & Gotthard, K. (2022). Local adaptation of life cycles in a butterfly is associated with variation in several circadian clock genes. Molecular Ecology, 31(5), 1461–1475. 10.1111/mec.16331

Lindestad, O., Nylin, S., Wheat, C. W., & Gotthard, K. (2024). Testing for variation in photoperiodic plasticity in a butterfly: Inconsistent effects of circadian genes between geographic scales. Ecology and Evolution, 14(7), e11713. 10.1002/ece3.11713

Lirakis, M., Nolte, V., & Schlötterer, C. (2022). Pool-GWAS on reproductive dormancy in *Drosophila simulans* suggests a polygenic architecture. G3: Genes, Genomes, Genetics, 12(3), jkac027. 10.1093/g3journal/jkac027

Low, K. H., Lim, C., Ko, H. W., & Edery, I. (2008). Natural variation in the splice site strength of a clock gene and species-specific thermal adaptation. Neuron, 60(6), 1054–1067. 10.1016/j.neuron.2008.10.048

Majercak, J., Sidote, D., Hardin, P. E., & Edery, I. (1999). How a circadian clock adapts to seasonal decreases in temperature and day length. Neuron, 24(1), 219–230. 10.1016/S0896-6273(00)80834-X

Manning, L., Sheth, J., Bridges, S., Saadin, A., Odinammadu, K., Andrew, D., Spencer, S., Montell, D., & Starz-Gaiano, M. (2017). A hormonal cue promotes timely follicle cell migration by modulating transcription profiles. Mechanisms of Development, 148, 56– 68. 10.1016/J.MOD.2017.06.003

Markow, T. A., & O’Grady, P. M. (2006). Drosophila: A guide to species identification and use. Academic Press.

Mathias, D., Jacky, L., Bradshaw, W. E., & Holzapfel, C. M. (2005). Geographic and developmental variation in expression of the circadian rhythm gene, *timeless*, in the pitcher-plant mosquito, *Wyeomyia smithii*. Journal of Insect Physiology, 51(6), 661–667. 10.1016/j.jinsphys.2005.03.011

Mayekar, H. V., Ramkumar, D. K., Garg, D., Nair, A., Khandelwal, A., Joshi, K., & Rajpurohit, S. (2022). Clinal variation as a tool to understand climate change. Frontiers in Physiology, 13, 880728. 10.3389/fphys.2022.880728

McDonald, M. J., Rosbash, M., & Emery, P. (2001). Wild-type circadian rhythmicity is dependent on closely spaced E boxes in the *Drosophila timeless* promoter. Molecular and Cellular Biology, 21(4), 1207–1217. 10.1128/mcb.21.4.1207-1217.2001

Minami, N., & Kimura, M. T. (1980). Geographical variation of photoperiodic adult diapause in *Drosophila auraria*. The Japanese Journal of Genetics, 55(5), 319–324. 10.1266/jjg.55.319

Minh, B. Q., Nguyen, M. A. T., & von Haeseler, A. (2013). Ultrafast approximation for phylogenetic bootstrap. Molecular Biology and Evolution, 30(5), 1188–1195. 10.1093/molbev/mst024

Minh, B. Q., Schmidt, H. A., Chernomor, O., Schrempf, D., Woodhams, M. D., von Haeseler, A., & Lanfear, R. (2020). IQ-TREE 2: new models and efficient methods for phylogenetic inference in the genomic era. Molecular Biology and Evolution, 37(5), 1530–1534. 10.1093/molbev/msaa015

Miyake, H., & Watada, M. (2007). Molecular phylogeny of the *Drosophila auraria* species complex and allied species of Japan based on nuclear and mitochondrial DNA sequences. Genes & Genetic Systems, 82(1), 77–88. 10.1266/ggs.82.77

Montelli, S., Mazzotta, G., Vanin, S., Caccin, L., Corrà, S., Pittà, C. D., Boothroyd, C., Green, E. W., Kyriacou, C. P., & Costa, R. (2015). *Period* and *timeless* mRNA splicing profiles under natural conditions in *Drosophila melanogaster*. Journal of Biological Rhythms, 30(3), 217–227. 10.1177/0748730415583575

Nanda, S., Defalco, T. J., Hui Yong Loh, S., Phochanukul, N., Camara, N., Doren, M. V., & Russell, S. (2009). Sox100B, a *Drosophila* group E Sox-domain gene, is required for somatic testis differentiation. Sexual Development, 3(1), 26–37. 10.1159/000200079

Nguyen, L. T., Schmidt, H. A., von Haeseler, A., & Minh, B. Q. (2015). IQ-TREE: A fast and effective stochastic algorithm for estimating maximum-likelihood phylogenies. Molecular Biology and Evolution, 32(1), 268–274. 10.1093/molbev/msu300

Oguma, Y., Jallon, J. M., Tomaru, M., Matsubayashi, H. (1996). Courtship behavior and sexual isolation between *Drosophila auraria* and *D. triauraria* in darkness and light, Journal of Evolutionary Biology, 9(6), 803–815. 10.1046/j.1420-9101.1996.9060803.x

Onuma, M., Kamimura, Y., & Sawamura, K. (2022). Genital coupling and copulatory wounding in the *Drosophila auraria* species complex (Diptera: Drosophilidae). Biological Journal of the Linnean Society, 135(1), 195–207. https://academic.oup.com/biolinnean/article/135/1/195/6381422

Ortiz, E. M. (2019) vcf2phylip v2.0: Convert a VCF matrix into several matrix formats for phylogenetic analysis. Zenodo. 10.5281/zenodo.2540861

Paradis, E., & Schliep, K. (2019). ape 5.0: an environment for modern phylogenetics and evolutionary analyses in R. Bioinformatics, 35(3), 526–528. 10.1093/bioinformatics/bty633

Pertea, M., Pertea, G. M., Antonescu, C. M., Chang, T. C., Mendell, J. T., & Salzberg, S. L. (2015). StringTie enables improved reconstruction of a transcriptome from RNA-seq reads. Nature Biotechnology, 33, 290–295. 10.1038/nbt.3122

Pittendrigh, C. S., & Takamura, T. (1987). Temperature dependence and evolutionary adjustment of critical night length in insect photoperiodism. Proceedings of the National Academy of Sciences of the United States of America, 84(20), 7169–7173. 10.1073/pnas.84.20.7169

Pruisscher, P., Nylin, S., Gotthard, K., & Wheat, C. W. (2018). Genetic variation underlying local adaptation of diapause induction along a cline in a butterfly. Molecular Ecology, 27, 3613–3626. 10.1111/mec.14829

Pruisscher, P., Nylin, S., Wheat, C. W., & Gotthard, K. (2021). A region of the sex chromosome associated with population differences in diapause induction contains highly divergent alleles at clock genes. Evolution, 75(2), 490–500. 10.1111/evo.14151

Purcell, S., Neale, B., Todd-Brown, K., Thomas, L., Ferreira, M. A. R., Bender, D., Maller, J., Sklar, P., De Bakker, P. I. W., Daly, M. J., & Sham, P. C. (2007). PLINK: A tool set for whole-genome association and population-based linkage analyses. The American Journal of Human Genetics, 81(3), 559–575. 10.1086/519795

R Core Team (2025). R: A Language and Environment for Statistical Computing. R Foundation for Statistical Computing, Vienna, Austria. Available at: https://www.R-project.org/

Rambaut, A. (2018). FigTree v1.4.4: Tree figure drawing tool. Institute of Evolutionary Biology, University of Edinburgh. Available at: http://tree.bio.ed.ac.uk/software/figtree/

Rice, G., David, J. R., Kamimura, Y., Masly, J. P., McGregor, A. P., Nagy, O., Noselli, S., Nunes, M. D. S., O’Grady, P., Sánchez-Herrero, E., Siegal, M. L., Toda, M. J., Rebeiz, M., Courtier-Orgogozo, V., & Yassin, A. (2019). A standardized nomenclature and atlas of the male terminalia of *Drosophila melanogaster*. Fly, 13(1–4), 51–64. 10.1080/19336934.2019.1653733

Sandrelli, F., Tauber, E., Pegoraro, M., Mazzotta, G., Cisotto, P., Landskron, J., Stanewsky, R., Piccin, A., Rosato, E., Zordan, M., Costa, R., & Kyriacou, C. P. (2007). A molecular basis for natural selection at the *timeless* locus in *Drosophila melanogaster*. Science, 316(5833), 1898–1900. 10.1126/science.1138426

Schliep, K. P. (2011). phangorn: phylogenetic analysis in R. Bioinformatics, 27(4), 592–593. 10.1093/bioinformatics/btq706

Schmidt, P. S., Zhu, C. T., Das, J., Batavia, M., Yang, L., & Eanes, W. F. (2008). An amino acid polymorphism in the *couch potato* gene forms the basis for climatic adaptation in *Drosophila melanogaster*. Proceedings of the National Academy of Sciences of the United States of America, 105(42), 16207–16211. 10.1073/pnas.0805485105

Schneider, C. A., Rasband, W. S., & Eliceiri, K. W. (2012). NIH Image to ImageJ: 25 years of image analysis. Nature Methods, 9, 671–675. 10.1038/nmeth.2089

Sehadova, H., Glaser, F. T., Gentile, C., Simoni, A., Giesecke, A., Albert, J. T., & Stanewsky, R. (2009). Temperature entrainment of *Drosophila*’s circadian clock involves the gene nocte and signaling from peripheral sensory tissues to the brain. Neuron, 64(2), 251– 266. 10.1016/j.neuron.2009.08.026

Shafer, O. T., Levine, J. D., Truman, J. W., & Hall, J. C. (2004). Flies by night: effects of changing day length on *Drosophila*’s circadian clock. Current Biology, 14(5), 424–432. 10.1016/j.cub.2004.02.038

Shakhmantsir, I., Nayak, S., Grant, G. R., & Sehgal, A. (2018). Spliceosome factors target *timeless* (*tim*) mRNA to control clock protein accumulation and circadian behavior in *Drosophila*. eLife, 7, e39821. 10.7554/eLife.39821

Shen, W., Sipos, B., & Zhao, L. (2024). SeqKit2: A Swiss army knife for sequence and alignment processing. iMeta, 3, e191. 10.1002/imt2.191

Solares, E. A., Chakraborty, M., Miller, D. E., Kalsow, S., Hall, K., Perera, A. G., Emerson, J. J., & Hawley, R. S. (2018). Rapid low-cost assembly of the *Drosophila melanogaster* reference genome using low-coverage, long-read sequencing. G3: Genes, Genomes, Genetics, 8(10), 3143–3154. 10.1534/g3.118.200162

Sprengelmeyer, Q. D., Mansourian, S., Lange, J. D., Matute, D. R., Cooper, B. S., Jirle, E. V., Stensmyr, M. C., & Pool, J. E. (2020). Recurrent collection of *Drosophila melanogaster* from wild African environments and genomic insights into species history. Molecular Biology and Evolution, 37(3), 627–638. 10.1093/molbev/msz271

Suzuki, T., Kawai, T., Takemura, S., Nishiwaki, M., Suzuki, T., Nakamura, K., Ishiguro, S., & Higashiyama, T. (2018). Development of the Mitsucal computer system to identify causal mutation with a high-throughput sequencer. Plant Reproduction, 31, 117–128. 10.1007/S00497-018-0331-8

Tapanainen, R., Parker, D. J., & Kankare, M. (2018). Photosensitive alternative splicing of the circadian clock gene *timeless* is population specific in a cold-adapted fly, *Drosophila montana*. G3: Genes, Genomes, Genetics, 8(4), 1291–1297. 10.1534/g3.118.200050

Tauber, E., Zordan, M., Sandrelli, F., Pegoraro, M., Osterwalder, N., Breda, C., Daga, A., Selmin, A., Monger, K., Benna, C., Rosato, E., Kyriacou, C. P., & Costa, R. (2007). Natural selection favors a newly derived *timeless* allele in *Drosophila melanogaster*. Science, 316(5833), 1895–1898. 10.1126/science.1138412

Throckmorton, L. H. (1975). The phylogeny, ecology, and geography of *Drosophila*. In R.C. King (Ed.), Handbook of Genetics Vol. 3 (pp. 421–469). Plenum Press.

Trivers, R. L. (1972). Parental investment and sexual selection. In B. Campbell (Ed.), Sexual selection and the descent of man, 1871-1971 (pp. 136–179), Aldine.

Tyukmaeva, V., Lankinen, P., Kinnunen, J., Kauranen, H., & Hoikkala, A. (2020). Latitudinal clines in the timing and temperature-sensitivity of photoperiodic reproductive diapause in Drosophila montana. Ecography, 43, 759–768. 10.1111/ecog.04892

Vaze, K. M., Manoli, G., & Helfrich-Förster, C. (2024). *Drosophila ezoana* uses morning and evening oscillators to adjust its rhythmic activity to different daylengths but only the morning oscillator to measure night length for photoperiodic responses. Journal of Comparative Physiology A, 210, 535–548. 10.1007/s00359-023-01646-6

Watada, M., Matsumoto, M., Kondo, M., & Kimura, M. T. (2011). Taxonomic study of the *Drosophila auraria* species complex (Diptera: Drosophilidae) with description of a new species. Entomological Science, 14, 392–398. 10.1111/j.1479-8298.2011.00461.x

Weir, B. S., & Cockerham, C. C. (1984). Estimating F-statistics for the analysis of population structure. Evolution, 38(6), 1358–1370. 10.2307/2408641

Whitlock, M. C., & McCauley, D. E. (1999). Indirect measures of gene flow and migration: *F*_ST_≠1/(4Nm+1). Heredity, 82, 117–125. 10.1038/sj.hdy.6884960

Wickham, H. (2016). ggplot2: Elegant graphics for data analysis. Springer. 10.1007/978-3-319-24277-4

Willing, E. M., Dreyer, C., & van Oosterhout, C. (2012). Estimates of genetic differentiation measured by *F*_ST_ do not necessarily require large sample sizes when using many SNP markers. PLoS ONE, 7(8), e42649. 10.1371/journal.pone.0042649

Yamada, H., & Yamamoto, M. (2011). Association between circadian clock genes and diapause incidence in *Drosophila triauraria*. PLoS ONE, 6(12), e27493. 10.1371/journal.pone.0027493

Yu, Y., Huang, L. L., Xue, F. S., & Dopman, E. B. (2023). Partial reuse of circadian clock genes along parallel clines of diapause in two moth species. Molecular Ecology, 32, 3419–3439. 10.1111/mec.16940

Zheng, S., Wang, Y., Li, G., Qin, S., Dong, Z., Yang, X., Xu, X., Fang, G., Li, M., & Zhan, S. (2025). Functional polymorphism of CYCLE underlies the diapause variation in moths. Science, 388(6750), eado2129. 10.1126/science.ado2129

Zonato, V., Fedele, G., & Kyriacou, C. P. (2016). An intronic polymorphism in *couch potato* is not distributed clinally in European *Drosophila melanogaster* populations nor does it affect diapause inducibility. PLoS ONE, 11(9), e0162370. 10.1371/journal.pone.0162370

Zonato, V., Vanin, S., Costa, R., Tauber, E., & Kyriacou, C. P. (2018). Inverse European latitudinal cline at the *timeless* locus of *Drosophila melanogaster* reveals selection on a clock gene: population genetics of *ls-tim*. Journal of Biological Rhythms, 33(1), 15–23. 10.1177/0748730417742309

