## Supplemental_Information for "Geographic divergence and the genomic basis of reproductive diapause in *Drosophila triauraria*"

### Table of Contents:

| | |
|---|---|
| **Table S1** Summary of the whole-genome sequencing data quality and mapping statistics for each strain | Page 2 |
| **Table S2** List of *D. melanogaster* homologs identified in 5 kb genomic windows where *D. triauraria* high-latitude strains form a monophyletic cluster | Pages 3–4 |
| **Table S3** List of *D. melanogaster* homologs identified in 5 kb genomic windows where low-latitude strains form a monophyletic cluster | Pages 5–9 |
| **Figure S1** Accessory gland size immediately after eclosion in *D. triauraria* | Page 10 |
| **Figure S2** Monophyletic window analyses of the high- and low-latitudinal groups using 10, 20, and 100 kb windows | Pages 11–12 |
| **Figure S3** Sliding window analyses of $F_{ST}$ values between latitudinal strain groups in *D. triauraria* | Page 13 |
| **Figure S4** Venn diagrams comparing monophyletic windows with highly differentiated windows based on $F_{ST}$ | Page 14 |
| **Figure S5** Isoform group structures identified using 3'RACE and the amplified regions for RT-qPCR analysis | Page 15 |
| **Figure S6** Relative abundances of *tim* isoform groups in the heads of female *D. triauraria* from a high-latitude strain (ONMA20-3) | Page 16 |
| **Figure S7** Relative expression levels of *tim* isoform groups in the heads of female *D. triauraria* from high- mid- and low-latitude groups | Pages 17–18 |
| **Figure S8** The maximum likelihood trees of the whole *tim* region | Page 19 |

**Table S1 Summary of the whole-genome sequencing data quality and mapping statistics for each strain.**

| Species | Strain name | Latitude group | Raw reads (M) | Mapped reads [a] (M) | Mean read depth (× coverage) | Bases ≥ Q30 (%) |
|---|---|---|---|---|---|---|
| *D. triauraria* | SPR23-05 | High-latitude | 43.9 | 39.0 | 51.0 | 99.0 |
| *D. triauraria* | SPR23-09 | High-latitude | 49.1 | 44.2 | 57.5 | 99.0 |
| *D. triauraria* | TM15-38 | High-latitude | 56.4 | 51.0 | 66.2 | 99.0 |
| *D. triauraria* | TM15-57 | High-latitude | 52.0 | 45.9 | 60.5 | 99.0 |
| *D. triauraria* | TM15-67 | High-latitude | 44.2 | 38.3 | 48.4 | 99.0 |
| *D. triauraria* | TM15-92 | High-latitude | 46.0 | 41.5 | 55.1 | 98.9 |
| *D. triauraria* | ONMA20-3 | High-latitude | 75.3 | 50.4 | 62.0 | 92.1 |
| *D. triauraria* | ONM-15 | High-latitude | 45.5 | 39.3 | 38.8 | 84.4 |
| *D. triauraria* | TTR010 | Mid-latitude | 48.5 | 44.4 | 58.6 | 99.0 |
| *D. triauraria* | T544 | Mid-latitude | 46.7 | 41.8 | 53.9 | 99.0 |
| *D. triauraria* | TMU23-02 | Mid-latitude | 46.3 | 41.7 | 54.6 | 98.9 |
| *D. triauraria* | K445 | Mid-latitude | 63.6 | 54.1 | 54.5 | 87.3 |
| *D. triauraria* | Jou | Mid-latitude | 83.1 | 71.8 | 66.2 | 74.9 |
| *D. triauraria* | T748 | Mid-latitude | 18.7 | 15.7 | 20.9 | 94.8 |
| *D. triauraria* | Yak2 | Yakushima | 74.6 | 66.9 | 62.1 | 83.4 |
| *D. triauraria* | Yak3 | Yakushima | 65.0 | 58.4 | 49.7 | 82.6 |
| *D. triauraria* | YKS-MTK | Yakushima | 20.2 | 17.0 | 21.6 | 95.0 |
| *D. triauraria* | OEB12 | Low-latitude | 18.3 | 16.0 | 20.2 | 94.4 |
| *D. triauraria* | OKNG12-6 | Low-latitude | 22.2 | 19.6 | 25.2 | 94.6 |
| *D. triauraria* | KMJ1 | Low-latitude | 27.6 | 23.7 | 22.9 | 94.0 |
| *D. triauraria* | KMJD2 | Low-latitude | 20.2 | 17.8 | 31.2 | 94.9 |
| *D. auraria* | 14028-0471.01 | - | 42.5 | 32.5 | 28.4 | 94.9 |
| *D. auraria* | L5 | - | 18.0 | 14.9 | 19.5 | 95.9 |
| *D. auraria* | L8 | - | 13.1 | 11.0 | 14.0 | 95.4 |
| *D. auraria* | L11 | - | 19.3 | 15.7 | 20.0 | 93.5 |
| *D. biauraria* | SPR23-04 | - | 39.3 | 28.9 | 42.6 | 98.9 |
| *D. subauraria* | ONM-29 | - | 45.1 | 29.2 | 40.4 | 98.8 |

[a] Reads were mapped to the *D. triauraria* reference genome (GCA_014170255.2_RU_Dtri_1.1_genomic.fna).

**Table S2 List of *D. melanogaster* homologs identified in 5 kb genomic windows where *D. triauraria* high-latitude strains form a monophyletic cluster.**

| Window location[a] | | *D. melanogaster* homolog[b] |
|---|---|---|
| X | 29437001-29442000 | - |
| X | 29438001-29443000 | - |
| X | 29440001-29445000 | - |
| X | 29441001-29446000 | - |
| X | 29444001-29449000 | - |
| X | 29454001-29459000 | - |
| X | 29455001-29460000 | - |
| X | 29456001-29461000 | - |
| X | 29474001-29479000 | - |
| X | 29488001-29493000 | - |
| X | 29489001-29494000 | - |
| X | 29490001-29495000 | - |
| X | 29508001-29513000 | - |
| X | 29509001-29514000 | - |
| X | 29514001-29519000 | - |
| X | 29515001-29520000 | - |
| X | 29516001-29521000 | Graf |
| X | 29517001-29522000 | Graf |
| X | 29518001-29523000 | Graf |
| X | 29519001-29524000 | Graf |
| X | 29520001-29525000 | Graf |
| X | 29521001-29526000 | - |
| X | 29522001-29527000 | - |
| X | 29523001-29528000 | - |
| X | 29524001-29529000 | - |
| X | 29525001-29530000 | - |
| X | 29526001-29531000 | - |
| X | 29527001-29532000 | - |
| X | 29569001-29574000 | - |
| X | 29571001-29576000 | - |
| X | 29580001-29585000 | - |
| X | 29595001-29600000 | - |
| X | 29596001-29601000 | - |
| X | 29597001-29602000 | - |
| X | 29598001-29603000 | - |
| X | 29603001-29608000 | lncRNA:CR45464 |
| X | 30204001-30209000 | - |
| X | 30240001-30245000 | - |
| X | 30241001-30246000 | - |

(Table S2 continued from previous page)

| | | |
|---|---|---|
| X | 30242001-30247000 | - |
| X | 30243001-30248000 | - |
| X | 30245001-30250000 | - |
| X | 30246001-30251000 | - |
| X | 30247001-30252000 | - |
| X | 30248001-30253000 | - |
| X | 30249001-30254000 | - |
| X | 30250001-30255000 | - |
| X | 30251001-30256000 | - |
| X | 30252001-30257000 | - |
| X | 30253001-30258000 | - |
| X | 30254001-30259000 | - |
| X | 30279001-30284000 | - |
| X | 30280001-30285000 | - |
| X | 30281001-30286000 | - |
| X | 30282001-30287000 | - |
| 2L | 4336001-4341000 | tim, snoRNA:Or-CD16 |
| 2L | 4337001-4342000 | tim, snoRNA:Or-CD17 |
| 3R | 24379001-24384000 | Polr3C, CG5961 |

[a] Window locations are based on the *D. triauraria* reference genome (GCA_014170255.2_RU_Dtri_1.1_genomic.fna).
[b] Local BLASTN searches were conducted against the *D. melanogaster* transcript (GCF_000001215.4_Release_6_plus_ISO1_MT_rna.fna.gz ). Only hits with E-values < 0.01 are shown.

**Table S3 List of *D. melanogaster* homologs identified in 5 kb genomic windows where low-latitude strains form a monophyletic cluster.**

| Window location[a] | | *D. melanogaster* homolog[b] |
|---|---|---|
| X | 13816001-13821000 | CG15249 |
| X | 13817001-13822000 | CG15249 |
| X | 13818001-13823000 | - |
| X | 13819001-13824000 | - |
| 2L | 8896001-8901000 | - |
| 2L | 8898001-8903000 | - |
| 2L | 8899001-8904000 | - |
| 2L | 9360001-9365000 | - |
| 2L | 9361001-9366000 | - |
| 2L | 9362001-9367000 | - |
| 2L | 9744001-9749000 | - |
| 2L | 9745001-9750000 | lncRNA:CR44398 |
| 2L | 10373001-10378000 | lncRNA:CR45286 |
| 2L | 12457001-12462000 | - |
| 2L | 12458001-12463000 | - |
| 2L | 12459001-12464000 | - |
| 2L | 12460001-12465000 | - |
| 2L | 13147001-13152000 | CG9967 |
| 2L | 13149001-13154000 | CG9967, eys |
| 2L | 13264001-13269000 | lncRNA:CR45361, lncRNA:CR45360, mir-1, lncRNA:CR45359 |
| 2L | 13265001-13270000 | lncRNA:CR45361, lncRNA:CR45360, mir-1, lncRNA:CR45360 |
| 2L | 13430001-13435000 | - |
| 2L | 13437001-13442000 | Sfp38D |
| 2L | 13656001-13661000 | CG34109 |
| 2L | 13657001-13662000 | CG34109 |
| 2L | 13658001-13663000 | CG34109 |
| 2L | 13659001-13664000 | CG34109 |
| 2L | 13660001-13665000 | CG34109 |
| 2L | 13664001-13669000 | CG34109 |
| 2L | 14968001-14973000 | - |
| 2L | 14969001-14974000 | Ada1-1, Ada1-2 |
| 2L | 15193001-15198000 | - |
| 2L | 15194001-15199000 | - |
| 2L | 15195001-15200000 | - |
| 2L | 15196001-15201000 | - |
| 2L | 15257001-15262000 | OtopLb |
| 2L | 15294001-15299000 | - |

(Table S3 continued from previous page)

| | | |
|---|---|---|
| 2L | 15296001-15301000 | - |
| 2L | 15300001-15305000 | CG3277 |
| 2L | 15338001-15343000 | - |
| 2L | 15339001-15344000 | - |
| 2L | 16231001-16236000 | Sema1a |
| 2L | 17920001-17925000 | - |
| 2L | 17962001-17967000 | - |
| 2L | 18509001-18514000 | CG18557 |
| 2L | 18510001-18515000 | CG18557 |
| 2L | 18517001-18522000 | - |
| 2L | 18518001-18523000 | - |
| 2L | 18519001-18524000 | - |
| 2L | 18522001-18527000 | - |
| 2L | 18536001-18541000 | - |
| 2L | 18706001-18711000 | - |
| 2L | 18707001-18712000 | - |
| 2L | 18721001-18726000 | Edem2 |
| 2L | 18722001-18727000 | Edem2 |
| 2L | 19236001-19241000 | - |
| 2L | 19238001-19243000 | - |
| 2L | 21740001-21745000 | - |
| 2R | 571001-576000 | - |
| 2R | 572001-577000 | - |
| 2R | 573001-578000 | - |
| 2R | 574001-579000 | - |
| 2R | 575001-580000 | - |
| 2R | 576001-581000 | - |
| 2R | 577001-582000 | - |
| 2R | 578001-583000 | - |
| 2R | 579001-584000 | - |
| 2R | 580001-585000 | - |
| 2R | 586001-591000 | lncRNA:CR44781 |
| 2R | 589001-594000 | lncRNA:CR44782 |
| 2R | 591001-596000 | lncRNA:CR44782 |
| 2R | 592001-597000 | lncRNA:CR44782 |
| 2R | 593001-598000 | - |
| 2R | 596001-601000 | - |
| 2R | 599001-604000 | - |
| 2R | 600001-605000 | - |
| 2R | 601001-606000 | - |
| 2R | 602001-607000 | - |

(Table S3 continued from previous page)

| | | |
|---|---|---|
| 2R | 603001-608000 | - |
| 2R | 604001-609000 | lncRNA:CR44456 |
| 2R | 605001-610000 | lncRNA:CR44456 |
| 2R | 606001-611000 | Gpdh2, lncRNA:CR44456 |
| 2R | 4923001-4928000 | - |
| 2R | 4924001-4929000 | - |
| 2R | 5179001-5184000 | lncRNA:CR44448, lncRNA:TS18, lncRNA:CR44450 |
| 2R | 5180001-5185000 | lncRNA:TS18, lncRNA:CR44450 |
| 2R | 5181001-5186000 | lncRNA:CR44450 |
| 2R | 6204001-6209000 | Ir60a |
| 2R | 6205001-6210000 | - |
| 2R | 6331001-6336000 | CG3082 |
| 2R | 6333001-6338000 | CG3082 |
| 2R | 6337001-6342000 | DCTN3-p24 |
| 2R | 6338001-6343000 | DCTN3-p24 |
| 2R | 6339001-6344000 | DCTN3-p24 |
| 2R | 6342001-6347000 | - |
| 2R | 6344001-6349000 | - |
| 2R | 6345001-6350000 | - |
| 2R | 6515001-6520000 | nahoda |
| 2R | 6516001-6521000 | - |
| 2R | 6517001-6522000 | nahoda |
| 2R | 6518001-6523000 | nahoda |
| 2R | 6519001-6524000 | nahoda |
| 2R | 6520001-6525000 | nahoda |
| 2R | 6521001-6526000 | nahoda |
| 2R | 6522001-6527000 | nahoda |
| 2R | 6899001-6904000 | - |
| 2R | 6900001-6905000 | - |
| 2R | 6901001-6906000 | asRNA:CR43730, JhI-26 |
| 2R | 6902001-6907000 | asRNA:CR43730, JhI-26 |
| 2R | 7129001-7134000 | - |
| 2R | 7131001-7136000 | - |
| 2R | 9480001-9485000 | CG6484, asRNA:CR45275, CG14483, RhoGAP54D |
| 2R | 9862001-9867000 | - |
| 2R | 9863001-9868000 | - |
| 2R | 9864001-9869000 | - |
| 2R | 9865001-9870000 | - |
| 2R | 9866001-9871000 | - |
| 2R | 9867001-9872000 | - |
| 2R | 9868001-9873000 | - |

(Table S3 continued from previous page)

| | | |
|---|---|---|
| 2R | 11967001-11972000 | - |
| 2R | 11968001-11973000 | - |
| 2R | 11969001-11974000 | - |
| 2R | 11970001-11975000 | jeb |
| 2R | 11971001-11976000 | - |
| 2R | 11972001-11977000 | - |
| 2R | 11973001-11978000 | jeb |
| 2R | 12070001-12075000 | - |
| 2R | 12071001-12076000 | - |
| 2R | 12072001-12077000 | Cyp6a21 |
| 2R | 12073001-12078000 | - |
| 2R | 12074001-12079000 | Cyp6a8 |
| 2R | 12075001-12080000 | Cyp6a8 |
| 2R | 12076001-12081000 | Cyp6a8 |
| 2R | 12077001-12082000 | Cyp6a8, Cyp6a21 |
| 2R | 12078001-12083000 | Cyp6a8 |
| 2R | 12079001-12084000 | Cyp6a8 |
| 2R | 12766001-12771000 | - |
| 2R | 12767001-12772000 | - |
| 2R | 13493001-13498000 | - |
| 2R | 13494001-13499000 | - |
| 2R | 13495001-13500000 | polyph |
| 3R | 11667001-11672000 | Dscam3 |
| 3R | 11668001-11673000 | Dscam3 |
| 3R | 11669001-11674000 | - |
| 3R | 12487001-12492000 | - |
| 3R | 20789001-20794000 | - |
| 3R | 21944001-21949000 | - |
| 3R | 23054001-23059000 | DIP-gamma, CG11828 |
| 3R | 24072001-24077000 | - |
| 3R | 24073001-24078000 | - |
| 3R | 24683001-24688000 | Ccdc114 |
| 3R | 24684001-24689000 | Ccdc114 |
| 3R | 24685001-24690000 | Ccdc114 |
| 3R | 24686001-24691000 | Ccdc114, modSP |
| 3R | 24687001-24692000 | modSP |
| 3R | 25668001-25673000 | - |
| 3R | 25669001-25674000 | - |
| 3R | 25670001-25675000 | - |
| 3R | 25671001-25676000 | - |
| 3R | 25730001-25735000 | asRNA:CR46469, CG33337 |

(Table S3 continued from previous page)

| | | |
|---|---|---|
| 3R | 25834001-25839000 | eIF3d1, eIF3d2 |
| 3R | 25841001-25846000 | CG10254 |
| 3R | 25842001-25847000 | CG10254 |
| 3R | 25843001-25848000 | CG10254 |
| 3R | 25992001-25997000 | - |
| 3R | 25993001-25998000 | - |
| 3R | 26093001-26098000 | CG10000 |
| 3R | 26327001-26332000 | mesh |
| 3R | 26328001-26333000 | mesh |
| 3R | 26383001-26388000 | - |
| 3R | 26384001-26389000 | - |
| 3R | 26385001-26390000 | - |
| 3R | 26386001-26391000 | - |
| 3R | 26387001-26392000 | - |
| 3R | 26451001-26456000 | - |
| 3R | 26452001-26457000 | - |
| 3R | 26453001-26458000 | - |
| 3R | 26454001-26459000 | - |
| 3R | 26455001-26460000 | - |
| 3R | 26456001-26461000 | Sox100B |
| 3R | 26457001-26462000 | - |
| 3R | 27512001-27517000 | RpS7 |
| 3R | 32133001-32138000 | - |
| 3R | 32149001-32154000 | - |
| 3R | 33382001-33387000 | - |
| 3R | 36261001-36266000 | SP1029, CG31445, Psa |
| 3R | 36262001-36267000 | SP1029, CG31446, Psa |
| 3R | 36263001-36268000 | SP1029, CG31447, Psa |

[a] Window locations are based on the D. triauraria reference genome (GCA_014170255.2_RU_Dtri_1.1_genomic.fna).
[b] Local BLASTN searches were conducted against the *D. melanogaster* transcript (GCF_000001215.4_Release_6_plus_ISO1_MT_rna.fna.gz ). Only hits with E-values < 0.01 are shown.

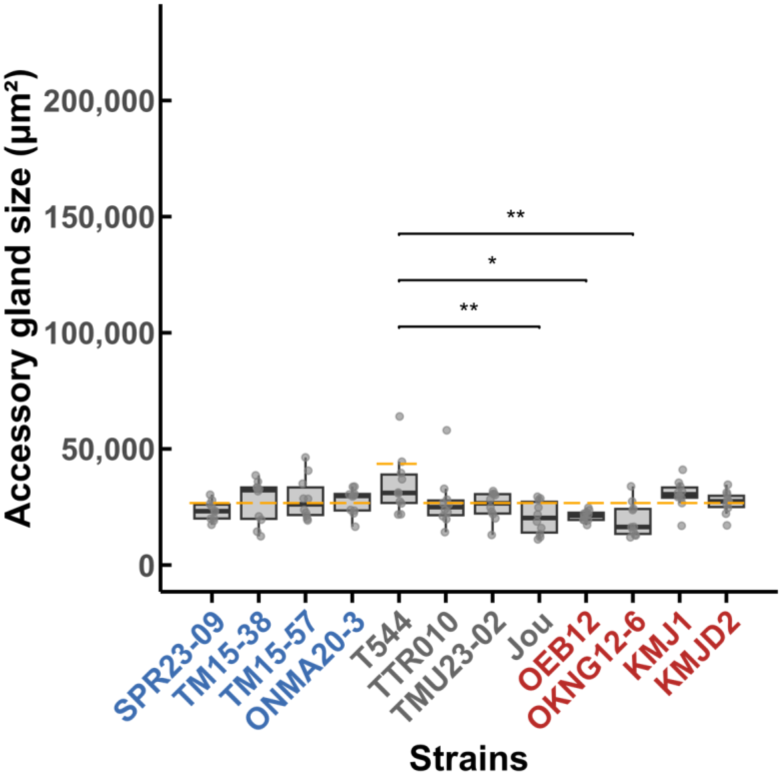


**FIGURE S1 Accessory gland size immediately after eclosion in *D. triauraria***
Accessory gland size within 6 h after eclosion (day 0), measured by area after mounting on the slides. $N = 10$ per strain. * $p<0.05$, ** $p<0.01$ by Tukey HSD test following ANOVA. Orange dashed lines indicate the upper limit of the 95% CI calculated within strain for T544, and for the remaining strains, from the pooled data excluding T544. The separate calculation for T544 was performed due to its significant difference from other strains. These upper-limit thresholds were used in Figure 4. Colors of the strain names represent their latitudinal origin: blue, grey, and red indicate high-latitude, mid-latitude, and low-latitude regions, respectively.

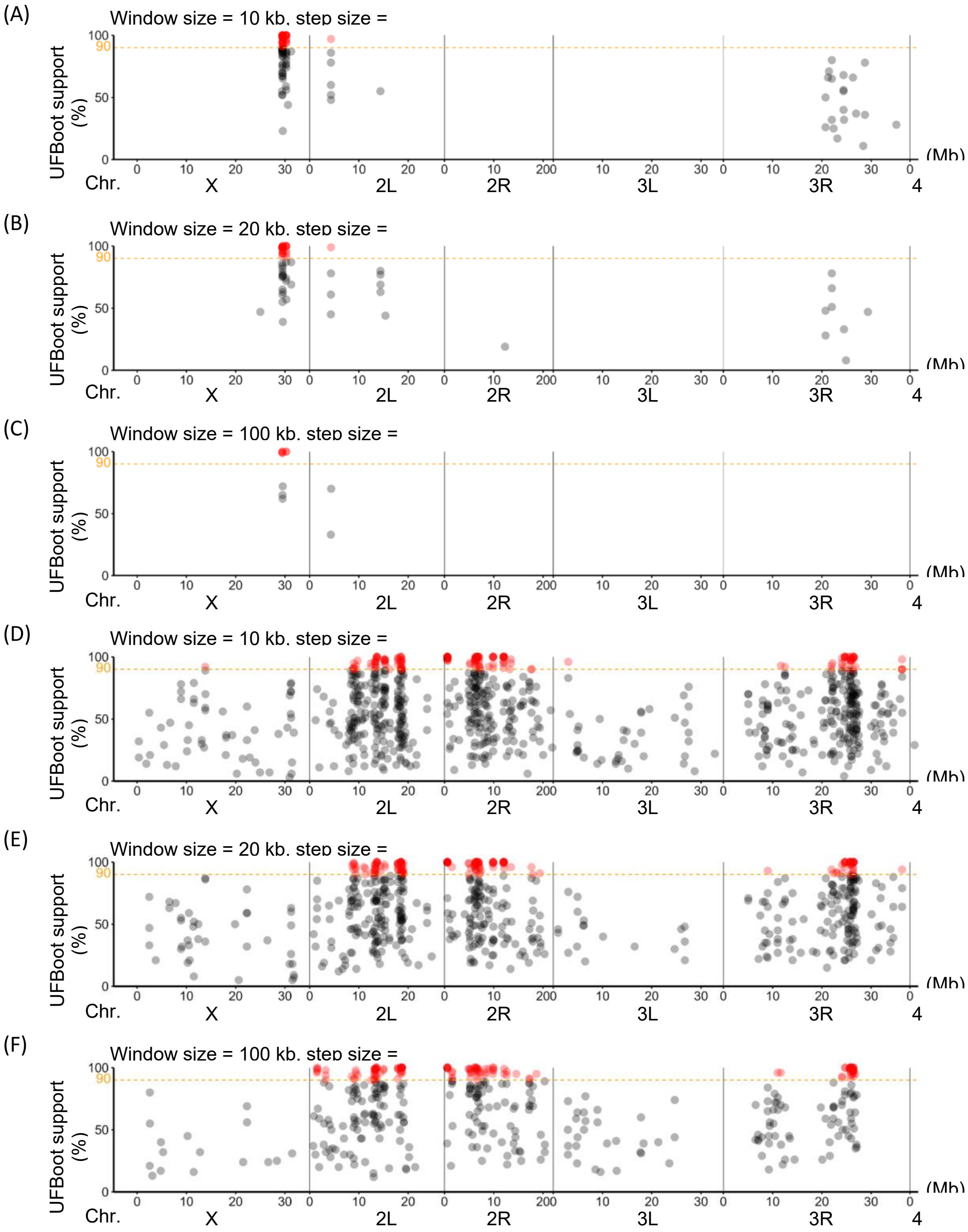
(A)
Window size = 10 kb, step size =
(B)
Window size = 20 kb, step size =
(C)
Window size = 100 kb, step size =
(D)
Window size = 10 kb, step size =
(E)
Window size = 20 kb, step size =
(F)
Window size = 100 kb, step size =
UFBoot support (%)
100
90
50
0
(Mb)
Chr.
0 10 20 30 0 10 20 0 10 200 10 20 30 0 10 20 30 0
X
2L
2R
3L
3R
4

(Figure S2)

**FIGURE S2 Monophyletic window analyses of the high- and low-latitudinal groups using 10, 20, and 100 kb windows**

Sliding-window profile of bootstrap (UFBoot) support (%) for monophyly of the high-latitude strain group (A–C) and the low-latitude strain group (D–F), based on maximum parsimony trees. (A, D) Window size = 10 kb; step size = 2 kb. (B, E) Window size = 20 kb; step size = 4 kb. (C, F) Window size = 100 kb; step size = 5 kb. Red dots mark the midpoints of windows where low-latitude strains formed monophyletic clusters with ≥ 90% bootstrap support (1,000 pseudo-replicates). The genome coordinates are based on the *D. triauraria* reference genome (GCA_014170255.2_RU_Dtri_1.1_genomic.fna).

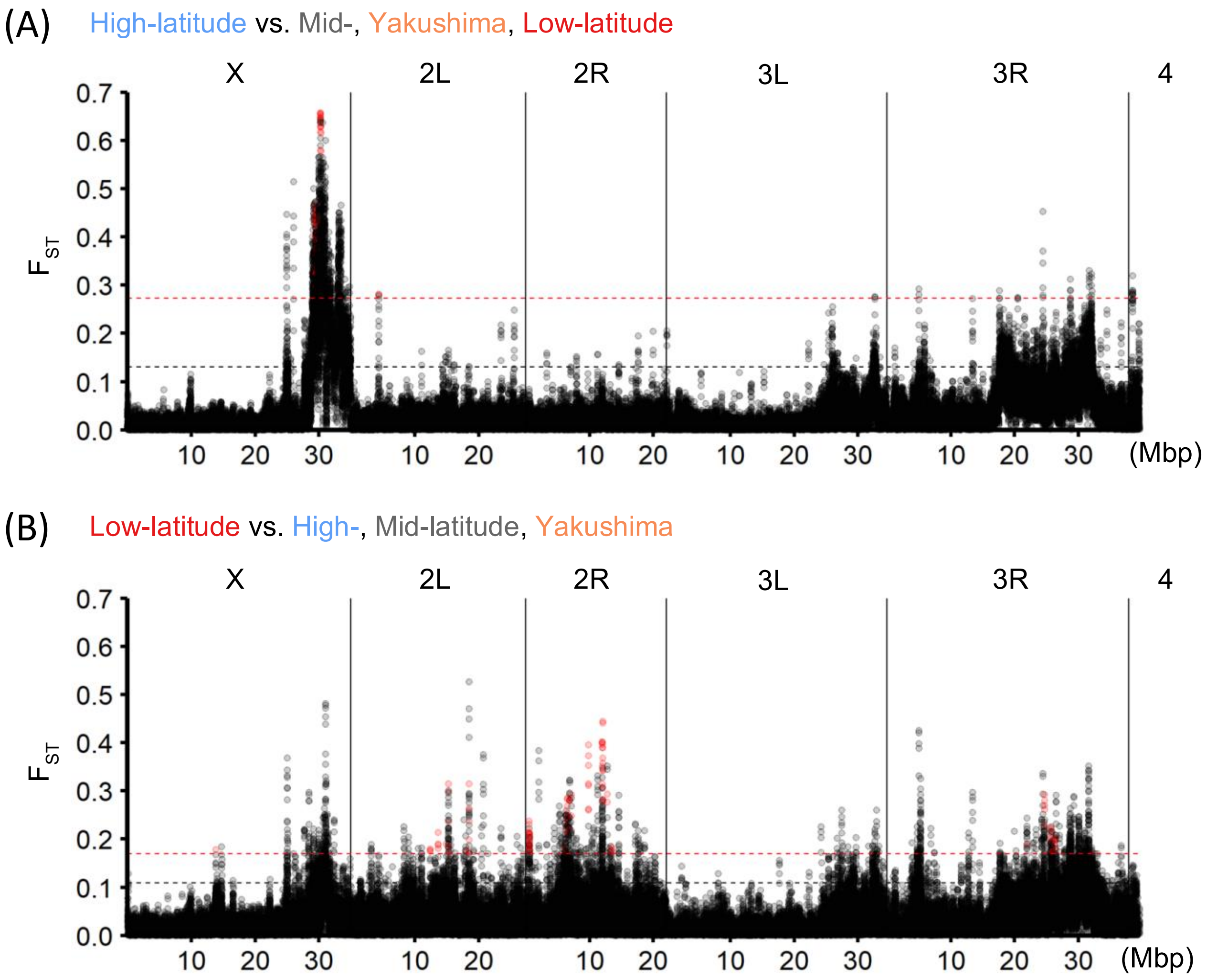


**FIGURE S3 Sliding window analyses of $F_{ST}$ values between latitudinal strain groups in *D. triauraria*.** (A) Pairwise weighted $F_{ST}$ values calculated between the high-latitude strain group and three other groups (mid-latidude, Yakushima, and low-latitude) for 5 kb windows with step size 1 kb. (B) Pairwise weighted $F_{ST}$ values calculated between the low-latitude strain group and three other groups (high-latidude, mid-latitude, and Yakushima) for 5 kb windows with step size 1 kb. Windows containing fewer than 100 variants and those with negative $F_{ST}$ values were excluded from the analysis. Black and red dotted lines indicate 90% and 95% percentiles, respectively. Red dots indicate monophyletic windows with ≥ 90% bootstrap support that were also within top 5% $F_{ST}$ windows. The genome coordinates are based on the *D. triauraria* reference genome (GCA_014170255.2_RU_Dtri_1.1_genomic.fna).

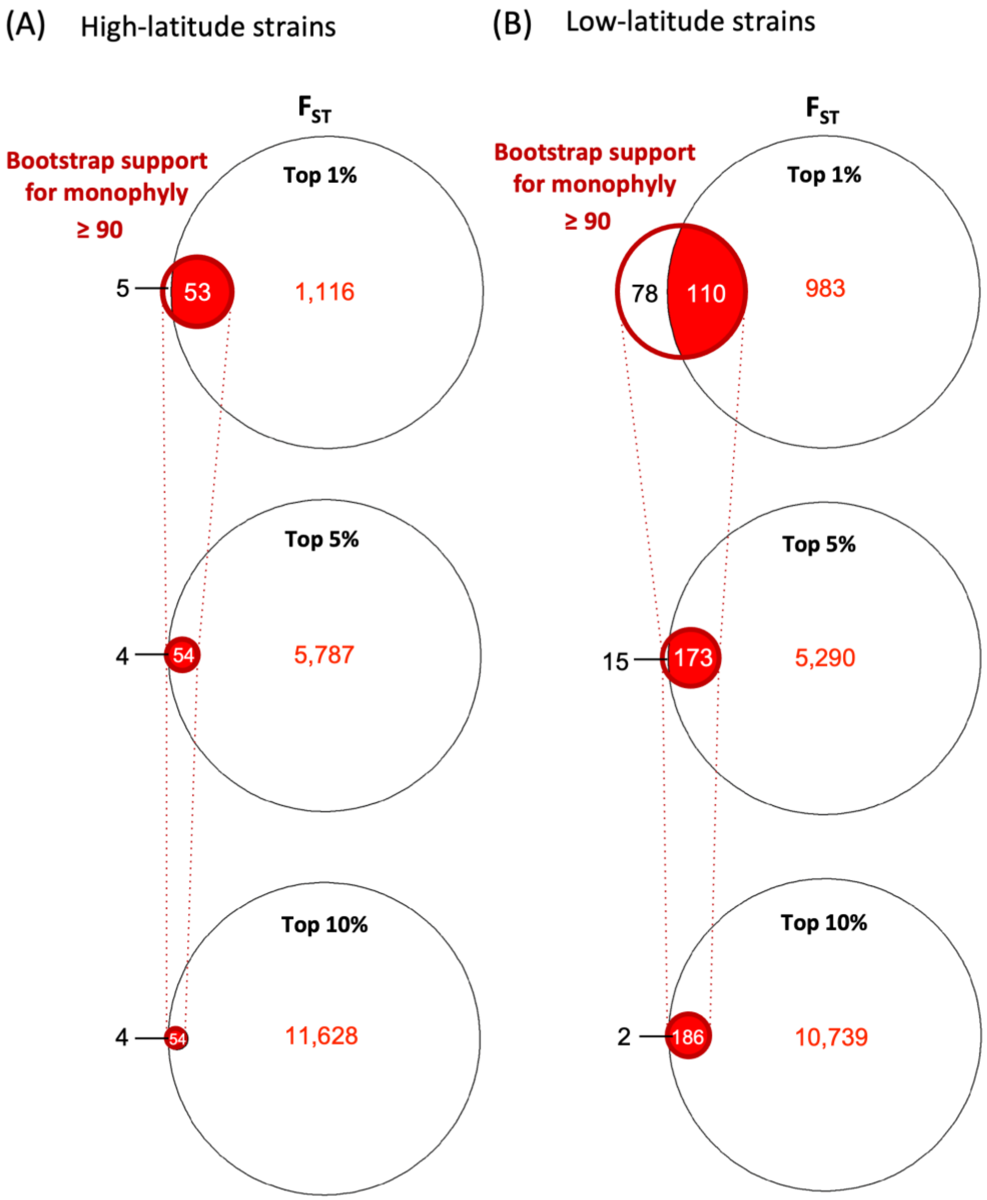


**FIGURE S4 Venn diagrams comparing monophyletic windows with highly differentiated windows based on $F_{ST}$.**

(A) Red-circled areas indicate windows with ≥90% bootstrap support for monophyly of the high-latitude strains. Black-circled areas represent windows within the top 1%, 5%, and 10% of $F_{ST}$ values between the high-latitude group and other latitudinal groups. Overlapping areas (red-filled) indicate genomic windows identified by both approaches. (B) Red-circled areas indicate windows with ≥90% bootstrap support for monophyly of the low-latitude strains. Black-circled areas represent windows within the top 1%, 5%, and 10% of $F_{ST}$ values between the low-latitude group and other latitudinal groups. Red-filled areas indicate genomic windows identified by both approaches.

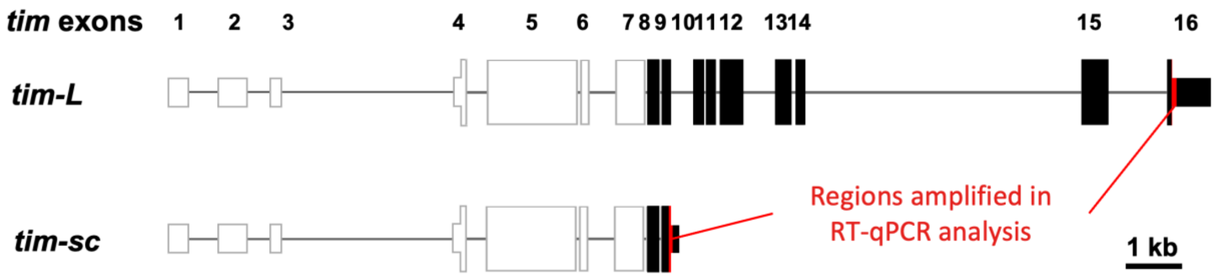


**FIGURE S5 Isoform group structures identified using 3'RACE and the amplified regions for RT-qPCR analysis.**

Sequences of *tim-L* isoform group (potentially including *tim-L*, *tim-cold* and other long isoforms) *and tim-sc* isoform group (potentially including *tim-sc* and other short isoforms) were identified in ONMA20-3 and OEB12 by 3'RACE. Black boxes indicate exons confirmed by 3'RACE. Gray boxes and connecting lines represent predicted exons and introns based on Hisat2, StringTie, and TransDecoder analyses. Red highlights the regions amplified in RT-qPCR analysis.

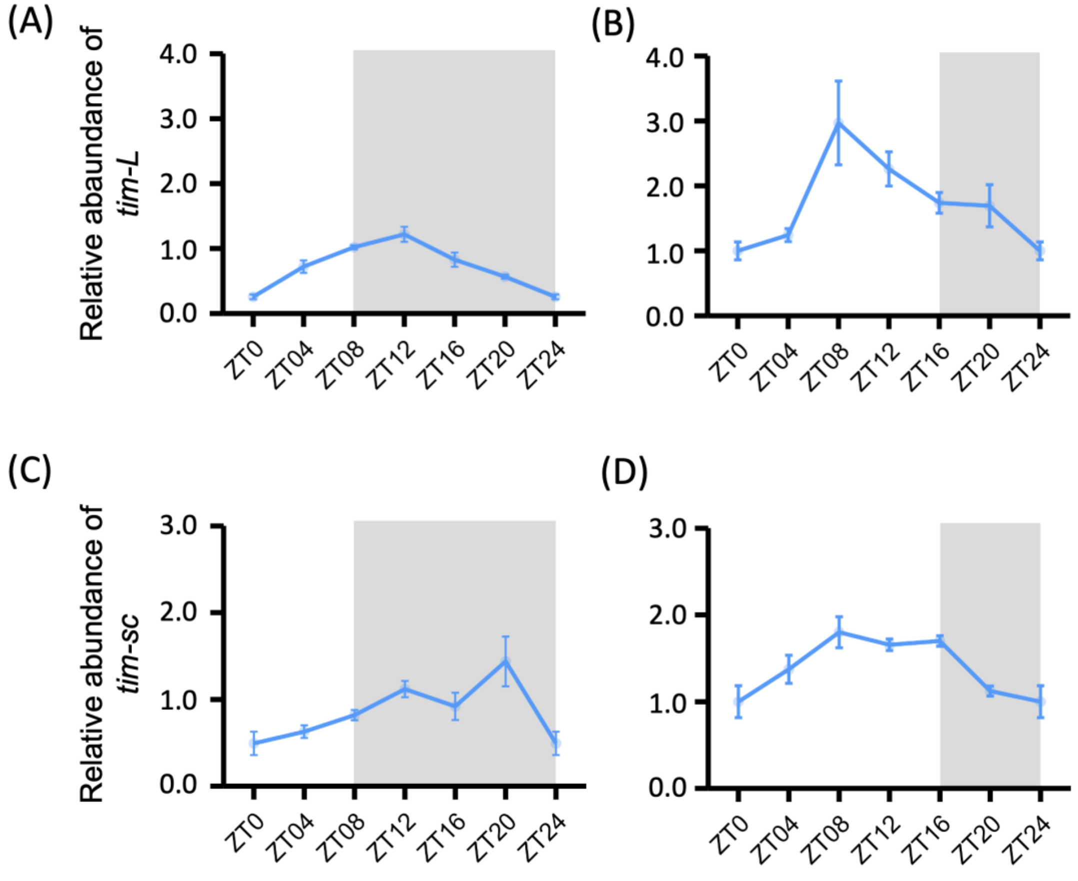


**FIGURE S6 Relative abundances of *tim* isoform groups in the heads of female *D. triauraria* from a high-latitude strain (ONMA20-3).**

(A) *tim-L* isoform group at six timepoints under 8L:16D. (B) *tim-L* isoform group at six timepoints under 16L:8D. (C) *tim-sc* isoform group at six timepoints under 8L:16D. (D) *tim-sc* isoform group at six timepoints under 16L:8D. Abundances of mRNA, measured by RT-qPCR, were normalized by *rp49* and standardized to the mean expression level at ZT0 under 16L:8D. Biological replicates: $N = 3$; technical replicates: $N = 2$.

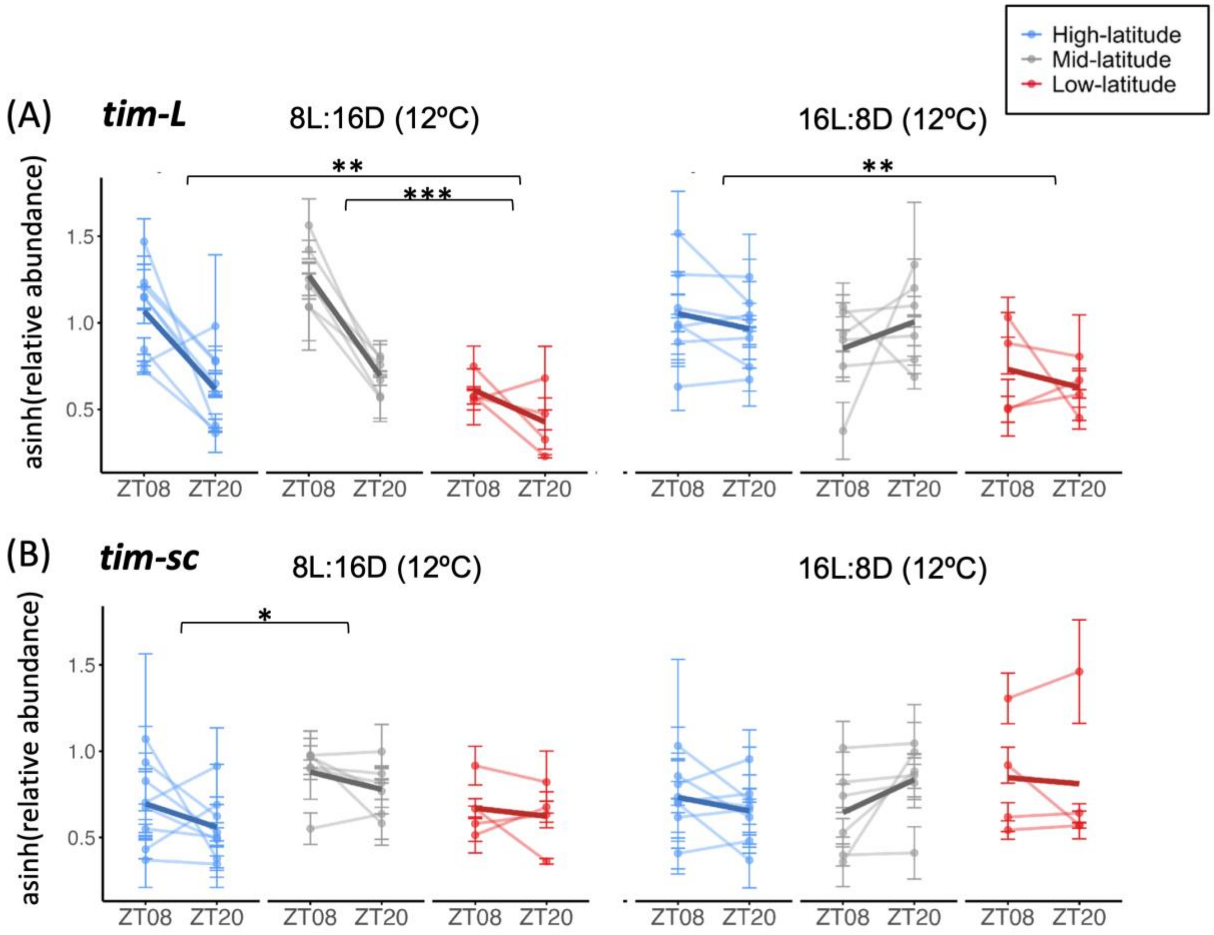


(C) **ANOVA summary table**

| Isoform group | Term | Df | 8L:16D | | 16L:8D | |
|---|---|---|---|---|---|---|
| | | | *F* value | *P* value | *F* value | *P* value |
| *tim-L* | Region | 2 | 13.108 | **< 0.0001** | 5.250 | **0.011** |
| | Time | 1 | 42.307 | **< 0.0001** | 0.019 | n.s. |
| | Region:Time | 2 | 2.337 | n.s. | 1.088 | n.s. |
| | Residuals | 30 | | | | |
| *tim-sc* | Region | 2 | 4.191 | **0.025** | 0.751 | n.s. |
| | Time | 1 | 2.668 | n.s. | 0.060 | n.s. |
| | Region:Time | 2 | 0.149 | n.s. | 1.015 | n.s. |
| | Residuals | 30 | | | | |

(Figure S7)

**FIGURE S7 Relative expression levels of *tim* isoform groups in the heads of female *D. triauraria* from high- mid- and low-latitude groups.**

(A) Relative expression of *tim-L* under 8L:16D and 16L:8D light/dark cycles at 12ºC, measured at ZT8 and ZT20. (B) Relative expression of *tim-sc* under 8L:16D and 16L:8D light/dark cycles at 12ºC, measured at ZT8 and ZT20. Expression levels were normalized to *rp49* and subjected to the hyperbolic arcsine transformation (asinh). Biological replicates: $N = 3$; technical replicates: $N = 2$. Asterisks above bars indicate significant pairwise differences by Tukey's HSD test: * $P < 0.05$, ** $P < 0.01$, *** $P < 0.001$. (C) Results of two-way ANOVA for *tim-L* and *tim-sc* isoform groups, testing the effects of region, time (ZT8 vs ZT20), and their interaction (region:time) under both light conditions. Significant effects ($P < 0.05$) are in bold. "n.s." indicates non-significant results.

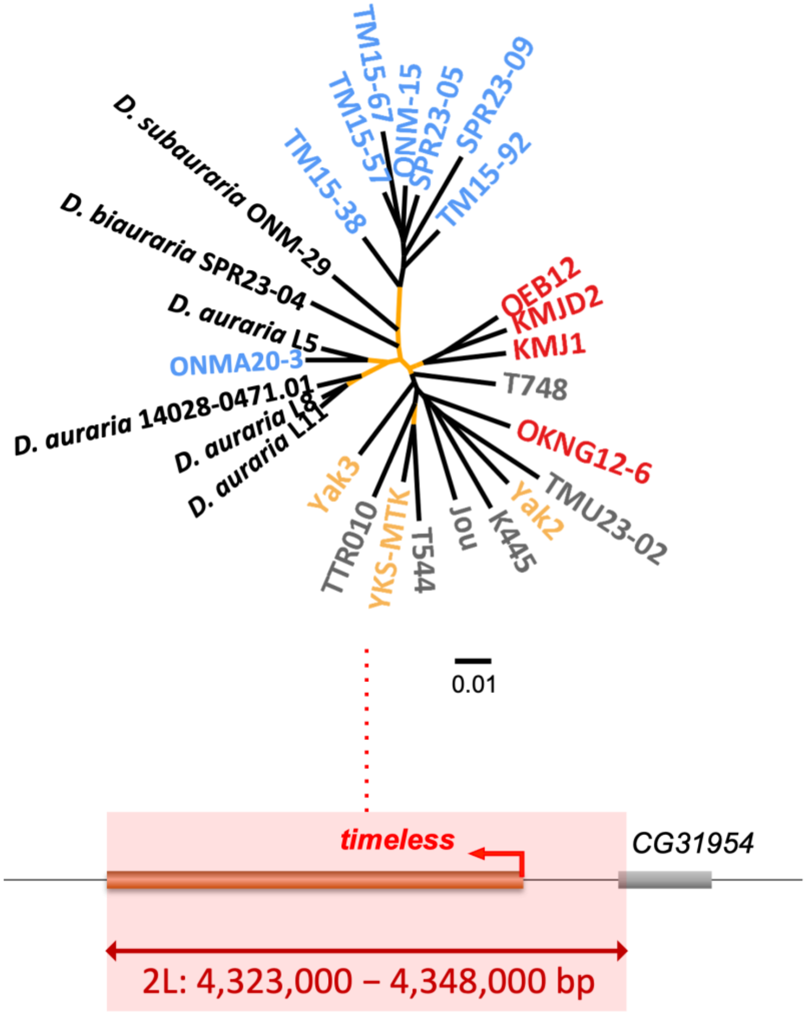


**FIGURE S8 The maximum likelihood trees of the whole *tim* region.**
Maximum likelihood tree using IQ-TREE with the GTR+G+I model constructed from the 25 kb region covering the whole *tim* gene (highlighted in pink). Nodes with UFBoot value ≥ 90%, estimated from 1,000 pseudo-replicates are highlighted in orange. The colors of *D. triauraria* strain names represent their latitudinal regions of origin: blue, grey, orange, and red indicate high-latitude, mid-latitude, Yakushima, and low-latitude regions, respectively.
